# Genome sequences of equine influenza A subtype H3N8 viruses by long read sequencing and functional characterization of the PB1-F2 virulence factor of A/equine/Paris/1/2018

**DOI:** 10.1101/2023.10.25.563963

**Authors:** Lena Kleij, Elise Bruder, Dorothée Raoux-Barbot, Nathalie Lejal, Quentin Nevers, Charlotte Deloizy, Bruno Da Costa, Loïc Legrand, Eric Barrey, Alexandre Chenal, Stéphane Pronost, Bernard Delmas, Sophie Dhorne-Pollet

## Abstract

Equine influenza virus (EIV) remains a persistent threat to equines, despite the availability of vaccines. Currently, strategies to monitor the virus and prevent any potential vaccine failure revolve around serological assays, RT‒qPCR amplification, and sequencing the viral hemagglutinin (HA) and neuraminidase (NA) genes. These approaches overlook the contribution of other viral proteins in driving virulence. This study assesses the potential of long-read nanopore sequencing for swift and precise sequencing of circulating equine influenza viruses. To this end, two French Florida Clade 1 strains, including the one circulating in winter 2018-2019 exhibiting more pronounced pathogenicity than usual, as well as the two currently used OIE-recommended vaccine strains, were sequenced. Our results demonstrated the reliability of this sequencing method in generating accurate sequences. Sequence analysis of HA revealed a subtle antigenic drift in the French EIV strains, with specific substitutions, such as T163I in A/equine/Paris/1/2018 and the N188T mutation in post-2015 strains; both substitutions were located in antigenic site B. Antigenic site E exhibited modifications in post-2018 strains, with the N63D substitution. Segment 2 sequencing also revealed that the A/equine/Paris/1/2018 strain encodes a longer variant of the PB1-F2 protein when compared to other Florida clade 1 strains (90 amino acids long versus 81 amino acids long). Further biological and biochemistry assays demonstrated that this PB1-F2 variant has enhanced abilities to abolish the mitochondrial membrane potential ΔΨm and permeabilize synthetic membranes. Altogether, our results highlight the interest in rapidly characterizing the complete genome of circulating strains with next-generation sequencing technologies to adapt vaccines and identify specific virulence markers of EIV.

## Introduction

Equine influenza (EI) is a highly contagious respiratory disease affecting horses, with significant economic repercussions on the global equine industry [1–4]. Its widespread transmission is facilitated by the international transport of horses, primarily for competition and breeding purposes [5, 6]. Common clinical manifestations of EI infection in naïve and unprotected animals include pyrexia, persistent cough, serous nasal discharge, dyspnea, muscle pain or weakness, lethargy, anorexia, and often complications arising from secondary bacterial infections [7, 8]. Although rarely fatal on its own, EI can lead to secondary bacterial infections in the respiratory tract and lungs, exacerbating the clinical condition of affected horses [4, 8]. Equine influenza virus (EIV), which is the causal agent of EI, is an influenza type A virus belonging to the *Orthomyxovirus* genus within the *Orthomyxoviridae* family. Currently, EI is known to be caused by only two primary virus subtypes: H3N8 and H7N7, with the latter remaining undetected since the 1970s [9]. The H3N8 subtype emerged in 1963 [10] in the Americas and has since spread globally, continuing to trigger epizootic events [2, 3, 11–13]. In the 1980s, H3N8 further diverged into American and Eurasian lineages [14]. The American lineage subsequently branched into the Kentucky, South American, and Florida sublineages [15]. The Florida sublineage underwent additional evolution in the early 2000s, resulting in two subtypes: Florida sublineage clade 1 (FC1) and Florida sublineage clade 2 (FC2) [16]. FC1 predominantly circulated in the Americas, while FC2 prevailed in Europe. However, this pattern shifted with the 2009 outbreak of an FC1 strain in Europe [17, 18]. Subsequently, EIV FC1 caused an outbreak of an unprecedented scale between late 2018 and 2019 in Europe [12, 19], with 53 outbreaks reported in France, 228 in the United Kingdom, and approximately 80 in Ireland [20, 21]. During the 2018 outbreak, vaccination coverage was substantial in France [20]. The vaccines used during these outbreaks are still considered effective by the World Organization for Animal Health Expert Surveillance Panel (OIE ESP) [20–22].

Currently, most diagnostic tests for EIV rely on detecting viral antigens or RT‒qPCR amplification of viral nucleic acids obtained from nasal swab samples. These two approaches have distinct trade-offs: antigen testing is swift but has limited sensitivity, while RT‒qPCR is more time-consuming but offers higher sensitivity. Moreover, data generated by these methods have limitations in providing insights into epidemiological links and vaccine effectiveness. In most cases, sequencing of the viral strains is performed posteriorly by Sanger sequencing using several segment-specific primers [23]. This technique is efficient but very time-consuming, and multiplexing is not possible. Therefore, there is a need to develop new diagnostic tools that combine speed, sensitivity, ability to detect coinfections, and comprehensive genome sequence information. Such methods are vital for effective health management strategies, including the identification of potential new virulence factors and the precise design of vaccines.

In this study, our objective was to genetically characterize the equine influenza H3N8 viruses circulating in France during the winters of 2009 and 2018 and, more specifically, to identify and characterize potential virulence determinants and antigenicity through whole-genome sequencing. Therefore, we used MinION long-read sequencing technology, which offers rapid sequencing and multiplex barcoding [24–27]. The viral strains A/equine/Beuvron-en-Auge/2/2009 and A/equine/Paris/1/2018, along with the OIE-recommended vaccine strains A/equine/Richmond/1/2007 and A/equine/South Africa/4/2003, were sequenced. Our results suggest that the accessory protein PB1-F2 may contribute to the virulence of the A/equine/Paris/1/2018 strain.

## Results

### Full-length genome sequencing strategy

The four selected EIV strains A/equine/Beuvron-en-Auge/2/2009 and A/equine/Paris/1/2018 as well as the OIE-recommended vaccine strains A/equine/Richmond/1/2007 and A/equine/South Africa/4/2003 were used to obtain complete amplicon sequences using the long-read sequencing technology developed by Oxford Nanopore Technology. The workflow used is described in **Fig. 1**. Direct RNA sequencing was carried out using the A/equine/South Africa/4/2003 strain to evaluate the relative sensitivity and accuracy of this approach (data not shown).

**Figure 1:**
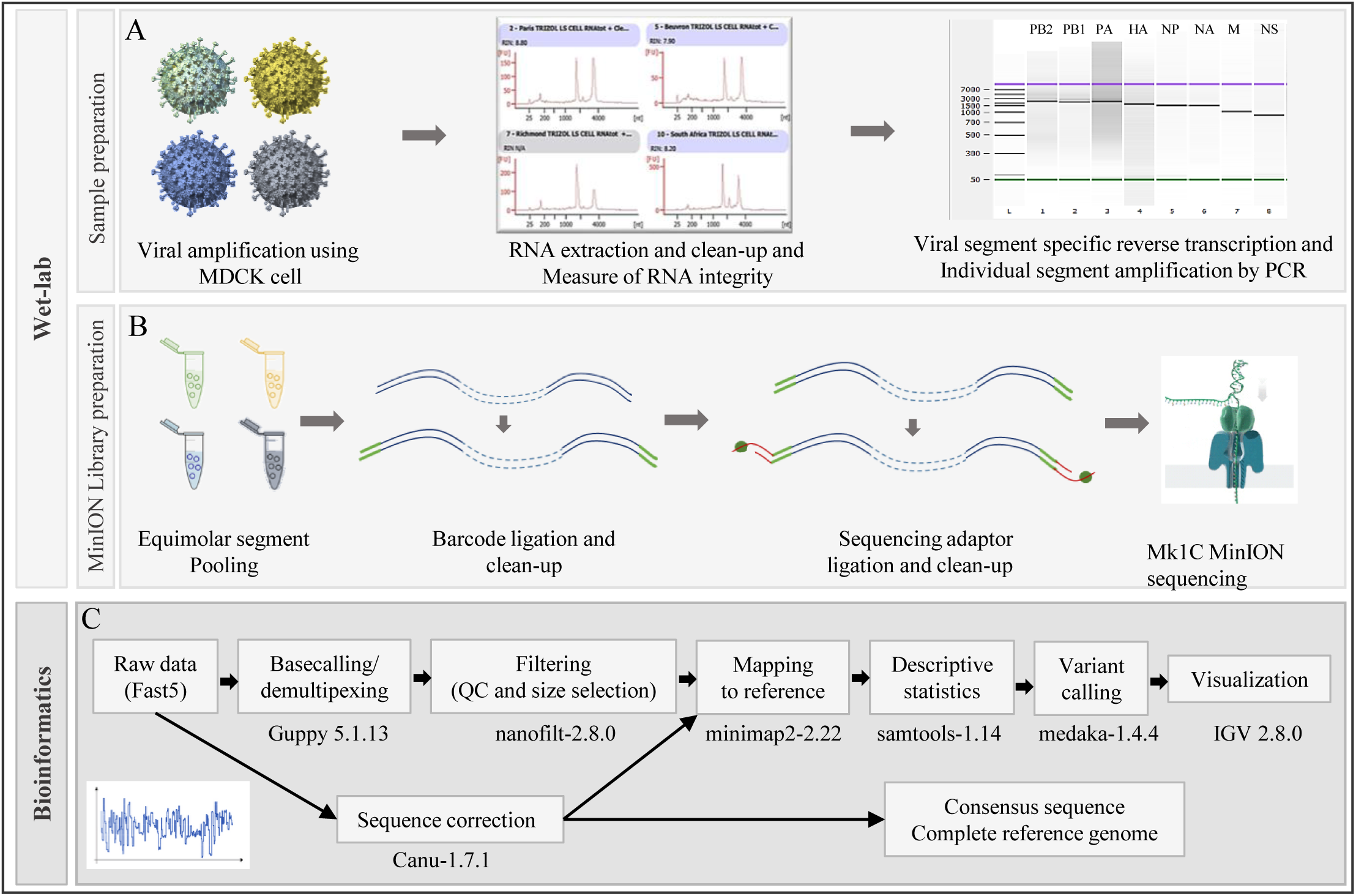
Schematic workflow implemented for long-read sequencing of equine influenza virus. The four equine influenza viruses A/equine/Beuvron-en-Auge/2/2009, A/equine/Paris/1/2018, and OIE recommended vaccine strains A/equine/South Africa/4/2003 and A/equine/Richmond/1/2007 were analyzed. (A) After viral amplification in the MDCK cell line and RNA extraction, the eight genomic segments were individually amplified by RT‒PCR. Amplified DNA products were controlled by capillary electrophoresis. (B) For each strain, the eight amplicons were pooled with equimolar ratios, and sequencing libraries were prepared and loaded on a flow cell. (C) The bioinformatics workflow used from raw data to consensus sequence construction. The reference strain is A/equine/Ohio/2005 (GenBank accession numbers: CY067323, CY067324, CY067325, CY067326, CY067327, CY067328, CY067329, CY067330).

After size and quality filtering, a mean of 235 222 reads per strain with 158 077 reads for A/equine/South Africa/4/2003, 173 495 reads for A/equine/Richmond/1/2007, 189 296 for A/equine/Paris/1/2018 and 420 018 reads for A/equine/Beuvron-en-Auge/2/2009 were produced (detailed sequencing statistics in **Table 1**). The average read length was 1291 bp for A/equine/South Africa/4/2003, 1132 bp for A/equine/Richmond/1/2007, 1215 bp for A/equine/Paris/1/2018 and 974 bp for A/equine/Beuvron-en-Auge/2/2009. The average quality (Phred score) for the four strains was Q=22. For the four strains, a mapping rate varying between 99.81% and 99.97% with full coverage of the eight influenza genome segments was obtained using the reference genome A/equine/Ohio/113461-1/2005.

**Table 1.** Sequencing statistics. Detailed sequencing statistics obtained after demultiplexing and size and quality filtering of the sequenced strains A/equine/South Africa/4/2003 (South Africa), A/equine/Richmond/1/2007 (Richmond), A/equine/Beuvron-en-Auge/2/2009 (Beuvron), and A/equine/Paris/1/2018 (Paris).

|  |  | South Africa | Richmond | Beuvron | Paris |
| --- | --- | --- | --- | --- | --- |
| **Total reads** |  | 515 999 | 519 851 | 976 418 | 582 633 |
| **Filtered reads**<br>(Q>10; Size>600 bp) | Reads number | 158 077 | 173 495 | 420 018 | 189 296 |
|  | Average length (nt) | 1 291 | 1 132 | 974 | 1 215 |
|  | Average quality | 22.5 | 22.6 | 22.1 | 22.6 |
| **Mapped reads** | Total reads mapped | 157 979 | 173 393 | 419 899 | 188 939 |
|  | % reads mapped | 99.94% | 99.94% | 99.97% | 99.81% |
|  | Segment1 (PB2) | 20 197 | 27 727 | 80 078 | 26 715 |
|  | Segment2 (PB1, PB1-F2) | 18 731 | 26 059 | 135 601 | 26 609 |
|  | Segment3 (PA) | 19 554 | 24 642 | 31 455 | 33 794 |
|  | Segment4 (HA) | 17 898 | 21 245 | 35 347 | 28 913 |
|  | Segment5 (NP) | 27 086 | 16 715 | 19 096 | 20 082 |
|  | Segment6 (NA) | 23 350 | 21 141 | 29 387 | 21 860 |
|  | Segment7 (M1-M2) | 26 425 | 35 844 | 54 494 | 35 756 |
|  | Segment8 (NS1, NEP) | 23 634 | 17 339 | 47 279 | 25 569 |

The nucleotide sequences of the viral genomes of the four strains were compared to those of A/equine/Ohio/113461-1/2005 (**Fig. 2, Supp Figs. 1 and 2**). No nucleotide discrepancies were observed between the genome sequence generated by amplicons and direct RNA sequencing of A/equine/South Africa/4/2003 (data not shown). A total of 538 substitutions for the four strains were detected. The A/equine/Paris/1/2018 genome exhibited a higher number of nucleotide substitutions (287 substitutions), particularly in the HA and NA segments, with 45 substitutions for each. Additionally, higher nucleotide sequence diversity was found in segments 1 and 3, encoding RNA-polymerase (FluPol) subunits PB2 and PA, respectively, with 53 and 49 substitutions among them and 32 and 31 being specific to A/equine/Paris/1/2018.

**Figure 2:**
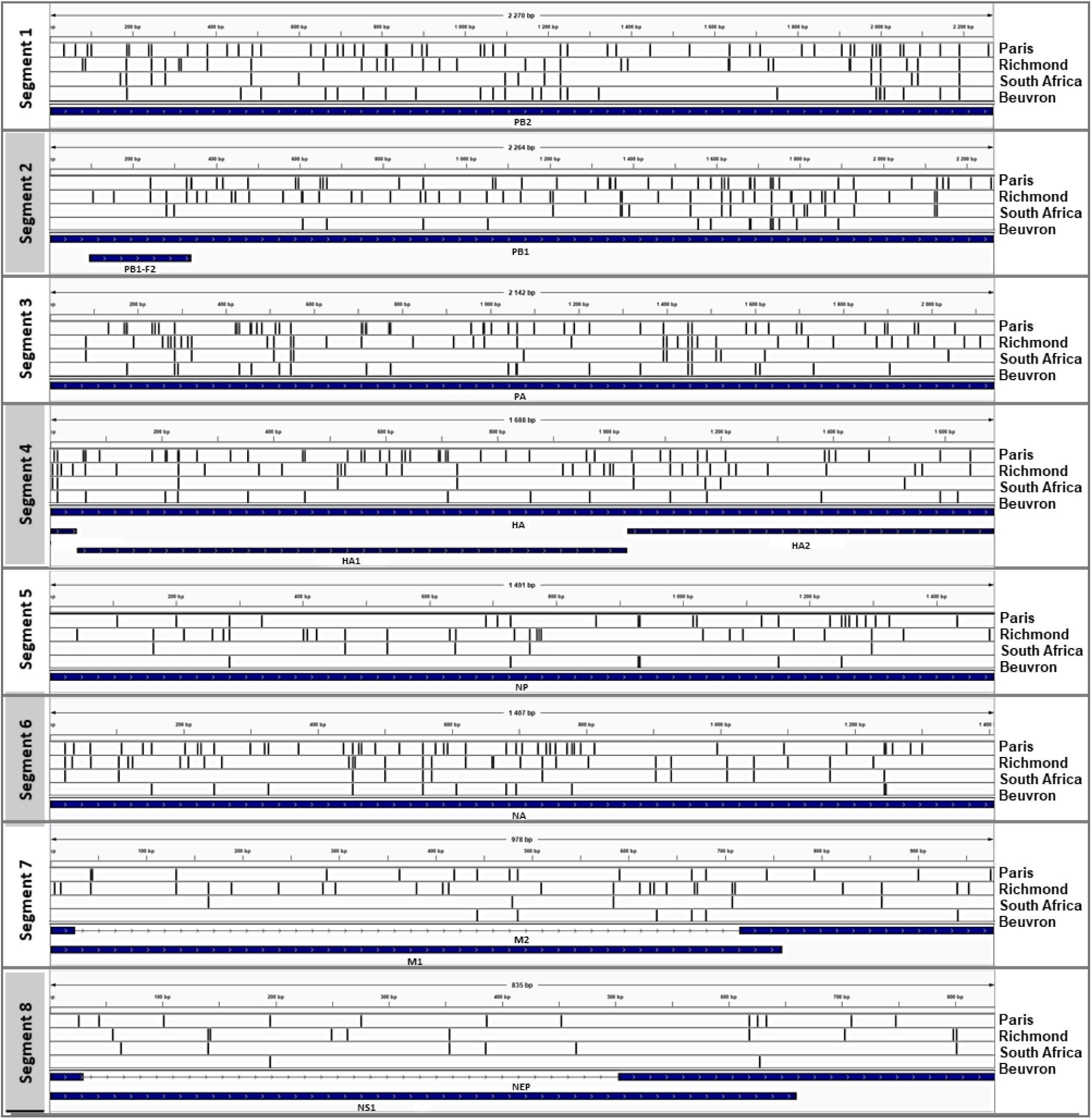
Nucleotidic variation patterns. This graphic extracted from Integrative Genomics Viewer [76] depicts variants as vertical bars along the x-axis for the different sequences shown on the y-axis. The four consensus genomic sequences of A/equine/Paris/1/2018 (Paris), A/equine/Richmond/1/2007 (Richmond), A/equine/South Africa/4/2003 (South Africa) and A/equine/Beuvron-en-Auge/2/2009 (Beuvron) are aligned to the reference (A/equine/Ohio/113461-1/2005 sequences) to visualize the variation patterns across the strains. The scale is indicated for each segment.

### Phylogenetic analyses

Individual phylogenetic trees were constructed for each of the eight segments, including sequences from the literature. The accession numbers of the selected sequences are presented in **Supp Table 1**. **Fig. 3** shows the analysis of complete HA and NA coding sequences. From 2011, the French isolates were present in both the FC1 and FC2 strains, with the A/equine/Paris/1/2018 HA segment exhibiting a higher phylogenetic distance from the vaccine strains. These observations for the HA gene were correlated with the complete NA sequence analysis. **Fig. 4A-D** shows the phylogenetic trees of the four segments encoding the components of the influenza ribonucleoprotein complex (with NP and FluPol subunits PA, PB1, and PB2). While all the phylogenetic trees correlate well with those of the HA and NA segments, the PA and PB1 subunits of A/equine/Saone-et-Loire/1/2015 exhibited a higher divergence than those of the other viruses, possibly reflecting the mark of a reassortment event with these two segments. In the same way, the phylogenetic trees carried out on segments 7 (M) and 8 (NS) correlated well with the ones described above, with exceptions in segment 7 of the two FC2 viruses (A/equine/Jouars/4/2006 and A/equine/Newmarket/5/2003) that appear to segregate with FC1 viruses and in segment 8 of two FC1 strains (A/equine/South Africa/4/2003 and A/equine/Ohio/1/2003) that group with FC2 viruses (**Fig. 4E-F**).

**Figure 3:**
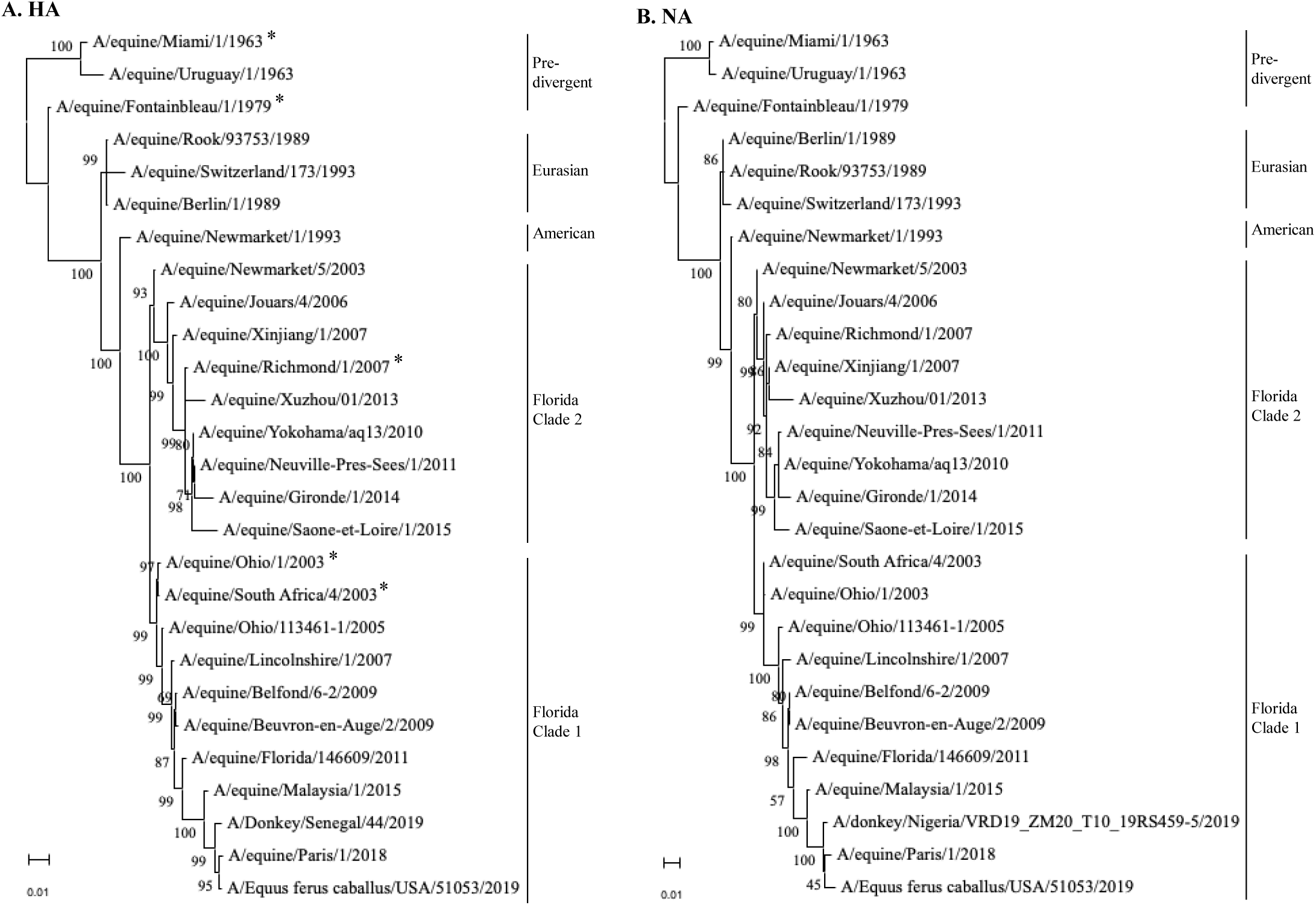
Phylogenetic analysis of the HA (A) and NA (B) nucleotide sequences for 27 EIV strains. The analysis includes representative strains of the main lineages, sublineages, and vaccine strains (*). Phylogenetic trees were created using the maximum likelihood method and Hasegawa-Kishino-Yano model with 1000 bootstraps.

**Figure 4:**
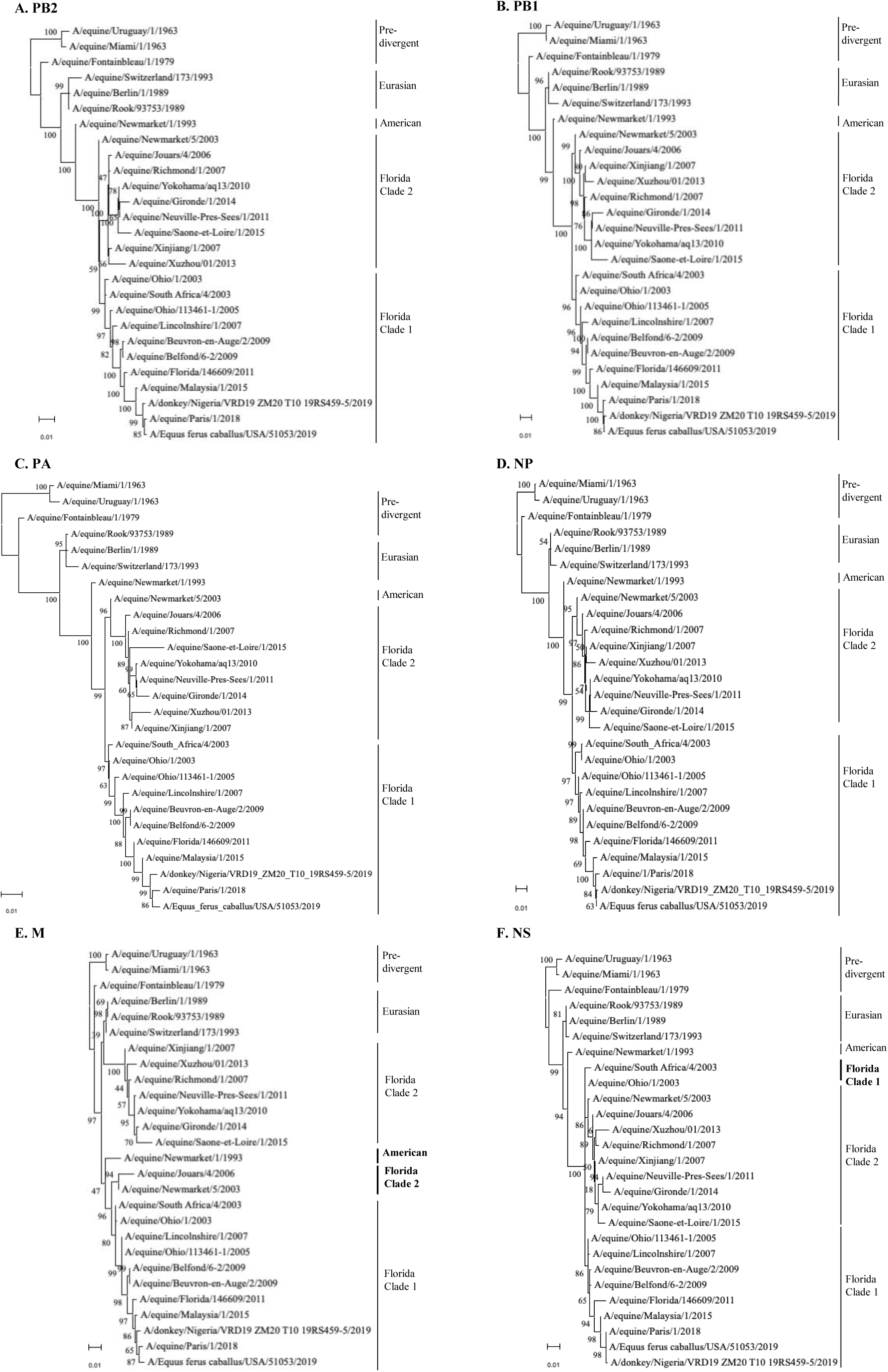
Phylogenetic analysis of the nucleotide sequences encoding PB2 (A), PB1 (B), PA (C), NP (D), M (E) and NS (F). The analysis includes representative strains of the main lineages, sublineages, and vaccine strains. Phylogenetic trees were created using the maximum likelihood method and Hasegawa-Kishino-Yano model with 1000 bootstraps.

### Analysis of HA amino acid alignment between circulating and ancestral viruses with vaccine strains

#### The antigenic sites

Five antigenic sites (A-E) have been previously defined on the hemagglutinin of influenza viruses of the H3 type (**Fig. 5 and Supp Fig. 3**), [28–31]). **Fig. 5A** shows a multiple alignment of amino acid sequences defining these antigenic sites on a selection of equine H3N8 viruses. **Fig. 5B** highlights the positions of the antigenic sites on the HA 3D structure. The recently circulating virus strains A/equine/Paris/1/2018 and A/equine/Beuvron-en-Auge/2/2009 were included in the analysis, with viruses belonging to FC1 and FC2 with representatives of French EIV strains and vaccine strains currently used in France (A/equine/Ohio/1/2003 and A/equine/Richmond/1/2007). Relatively high stability of the antigenic sites was observed for the FC1 and FC2 viruses over 40 years when compared to the two viruses isolated in 1963, A/equine/Miami/1/1963 and A/equine/Uruguay/1/1963. Among the 101 residues constituting the antigenic sites, only 19 and 18 substitutions were identified in A/equine/Paris/1/2018 and A/equine/Saone-et-Loire/1/2015, respectively. When compared with the currently used vaccine strains, only four substitutions (R62K, N63D, A138S and N188T) between FC1 circulating strains and A/equine/Ohio/1/2003 and three (A144T, T192K and Q197R) between FC2 strains A/equine/Saone-et-Loire/1/2015 and A/equine/Richmond/1/2007 were identified.

**Figure 5:**
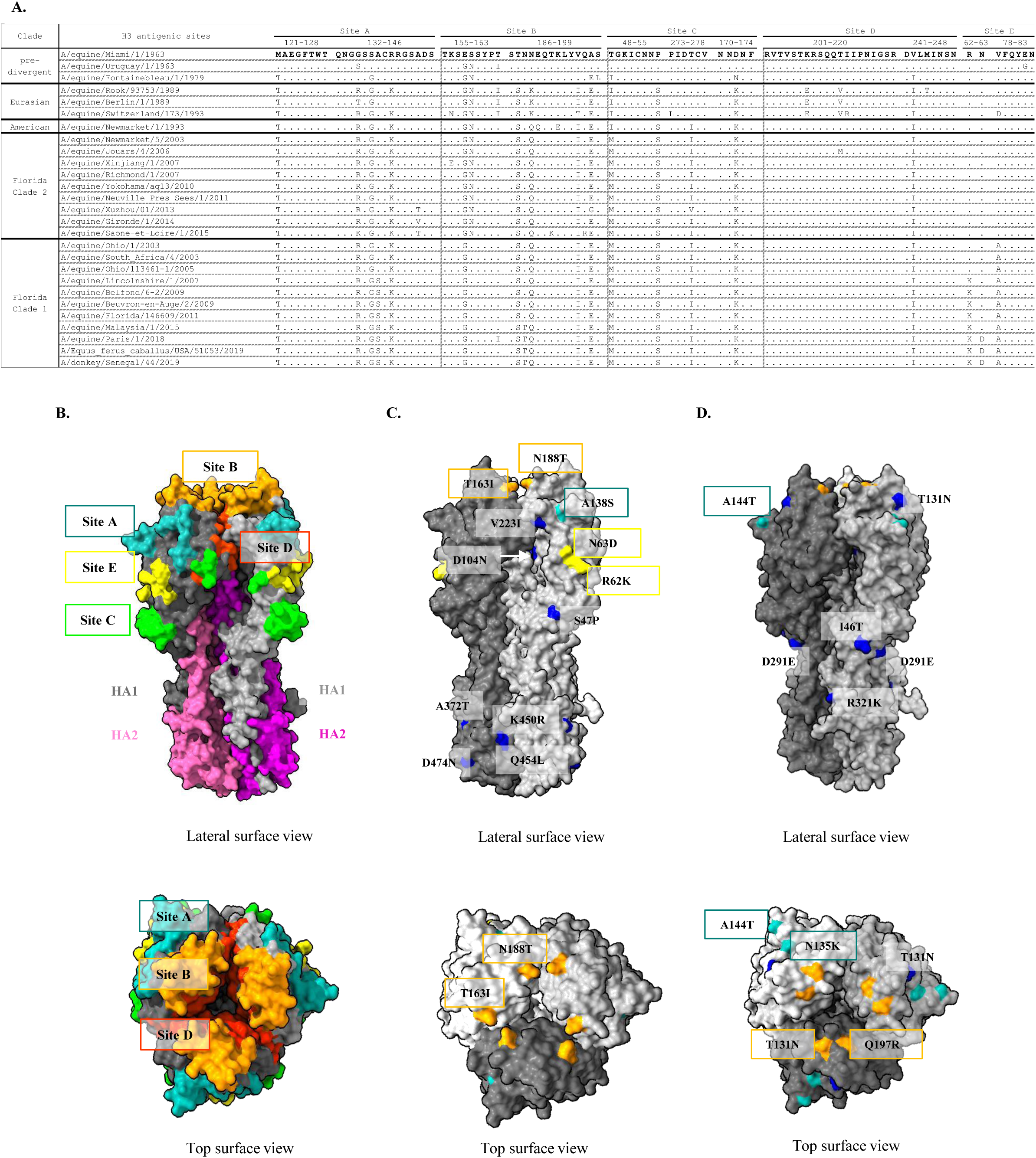
HA antigenic sites. (**A**) Amino acid alignments of the five antigenic sites A to E with HA sequences determined for French strains and other fully sequenced viral strains and compared with A/equine/Miami/1/1963. The antigenic sites defined for the human H3 influenza virus were used as a reference [28, 29, 31]. (**B**) Lateral and top views of the 3D structure of H3 hemagglutinin (PDB accession number: 4UO0) and location of its antigenic sites. While the HA2 domain (in pink and magenta) constitutes the stem, HA1 domains form the head of the HA bearing the antigenic sites. Antigenic sites are colored in cyan (site A), orange (site B), green (site C), red (site D), and yellow (site E). (**C**) Location of HA amino acid substitutions between the FC1 strains A/equine/Ohio/1/2003 and A/equine/Paris/1/2018. Amino acid changes are colored according to their positions in the corresponding antigenic sites (as in (B)) or in blue. (D) Location of HA amino acid substitutions between the FC2 strains A/equine/Richmond/1/2007 and A/equine/Saone-et-Loire/1/2015. Color patterning as in (C).

When restricting the analysis to three predivergent strains (with two of the 1963 years, the date of recognized emergence of H3N8 EIV), twelve amino acid substitutions occurred in the HA antigenic sites, several of them being conserved in subsequent clusters (T48I, M121T, G137G, E158G, S159N, T163I, A198E, and V242I). Others (E82G, G135S, D172N, and S199L) were not conserved among representatives of circulating strains of FC1 and FC2 when they diverged from 2003. Eurasian and American lineages (that emerged in the 1980s) displayed additional common substitutions (P55S, G135R/T, R140K, D172K, T187S, N189Q, and V196I) that were conserved in FC1 and FC2 circulating strains. Others (T48I, K156N, N189K, K207E, and T212V) were only represented in these two lineages. Among them, A/equine/Switzerland/173/1993 (Eurasian lineage) displayed additional specific substitutions (V78D, K156N, I213R, and P273L). A/equine/Newmarket/1/1993 (American lineage) also displayed a specific substitution (K193E). Concerning the FC1 and FC2 strains, T48M appeared to be the unique substitution marking these two sublineages. Other conserved substitutions (compared to the 1963 strains) were previously identified in the American lineage. The S159 variant was found only in the A/equine/Miami/1/1963 strain, and the V78A substitution is a hallmark of the FC1 strains when compared to other strains. As exemplified in **Fig. 5C**, several specific substitutions represented in different FC1 strains are R62K, N63D, A138S and N188T. For FC2 viruses, only one substitution in an antigenic site (A144T) was observed between the vaccine strain (A/equine/Richmond/1/2007) and the A/equine/Saone-et-Loire/1/2015 virus (**Fig. 5D**, [11]).

#### The receptor binding site

Because of the importance of receptor binding by HA in virus transmission and cross-species barriers, the analysis was extended to residues associated with binding to a2,3-linked receptors (**Supp. Fig. 3**). These residues are present on two loops on HA1, the 130-loop, the 220-loop, and the 190-helix [32, 33]. As expected, HA1 G225 and Q226 (220-loop), which are involved in receptor binding, are strictly conserved among all the strains analyzed. E190 and K193 are highly conserved (with two exceptions, E190Q and K193E in A/equine/Newmarket/1/1993). R135 and G137 (in the 130-loop and antigenic site A) exhibited full conservation in FC1 and FC2. Amino acid substitutions in the two loops were also identified in FC1 viruses (A138S and V223I).

#### The membrane fusion machinery

Two amino acid stretches in HA1 (a loop from residue 25 to 35) and HA2 (a-helix A between residues 367 and 384) constitute the fusion subdomain of HA that governs the fusion between cell and viral membranes. A single amino acid substitution, T30S, which was proposed to influence membrane fusion activity through local perturbation of the interactions between these two stretches [32], was identified in all FC1 and FC2 viruses. At position 379, a G379E substitution in several FC1 and FC2 viruses was observed. 3D structures of the HA of a Eurasian virus and an FC2 virus show that the glycine marks a break of the a-helix A [32], thus possibly modulating their fusion properties. The two HAs of the French strains A/equine/Paris/1/2018 and A/equine/Beuvron-en-Auge/2/2009 have a Gly at position 379.

Additional substitutions that are not involved in antigenic sites, receptor binding, or the fusion machinery are reported in **Supp. Table 2**.

### Analysis of NA amino acid alignment

Fourteen substitutions were identified between A/equine/Paris/1/2018 and the vaccine strain A/equine/Ohio/1/2003, seven in the stalk (A13T, N21S, V35A, G47E, T68I, I74M, R76K) and seven in the head (V147I, R252K, D258N, R260K, S337N, G416E and T434K) (**Supp. Table 2** and **Supp Fig. 4**). **Fig. 6** shows the substitutions exposed on the surface of the head of NA, one of them (V147I) located near the 150 loop of the active site [34].

**Figure 6:**
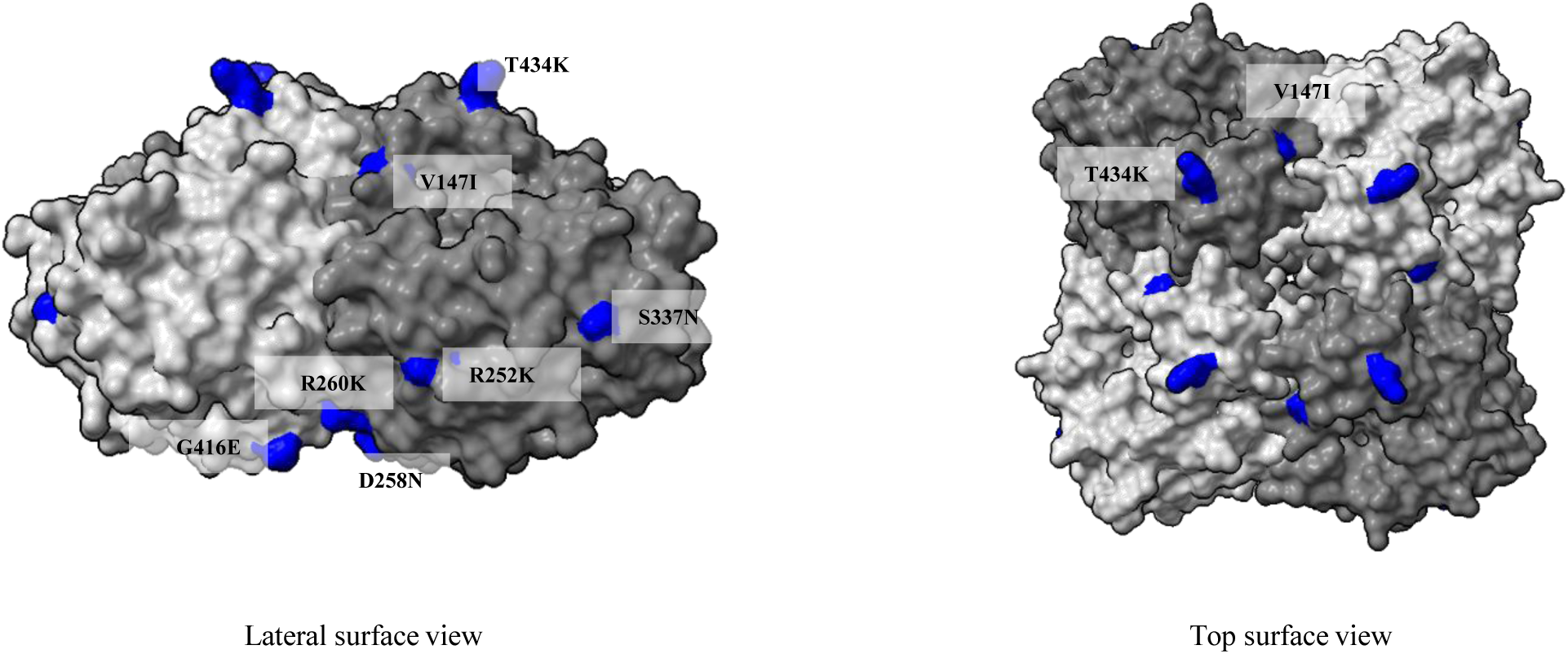
Positions of the amino acid substitutions on the surface of N8 between the FC1 vaccine strain A/equine/Ohio/1/2003 and the A/equine/Paris/1/2018 strain. Only the head of NA is represented. Amino acid changes are colored blue, and catalytic residues are colored yellow. The 3D structure template is the PDB accession number 2HT5.

### Comparison of the viral proteins of the replicative complex

The amino acid sequences of the FluPol subunits (PA, PB1 and PB2) and NP of the two FC1 strains A/equine/Paris/1/2018 and A/equine/Beuvron-en-Auge/2/2009 were compared with A/equine/Ohio/1/2003 and A/equine/Richmond/1/2007, the two OIE-recommended vaccine strains representing FC1 and FC2, respectively (**Fig. 7**). A greater number of changes in the EIV strain A/equine/Paris/1/2018 were identified in comparison to A/equine/Ohio/1/2003. This strain from 2018 possesses eight amino acid substitutions in PA, one in PB1, nine in PB2, and one in NP with A/equine/Ohio/1/2003. Some substitutions were also identified in A/equine/Beuvron-en-Auge/2/2009, such as in PB2 I63V, I398V, V667 and V686I, in PB1 F94L, R584Q and K621R and in PA E237K and T354I. Twenty-two substitutions between these two strains on the FluPol subunits and NP were also identified, exemplifying the continuous accumulation of substitutions between 2009 and 2018 in FC1 strains. Twenty-one substitutions between the two vaccine strains (isolated in 2003 and 2007) and three between A/equine/Ohio/1/2003 and A/equine/South Africa/4/2003 were also observed.

**Figure 7:**
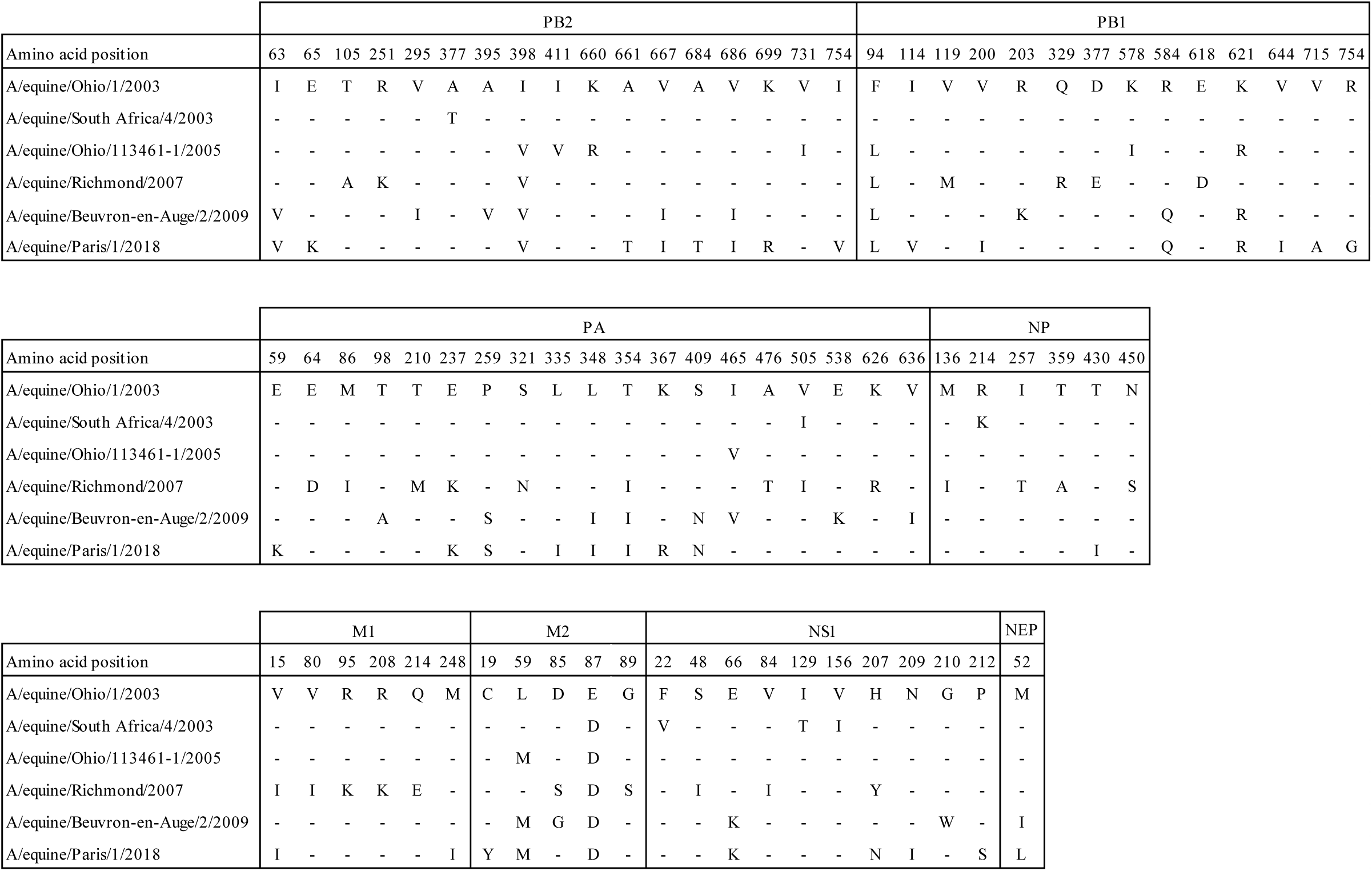
Amino acid sequence comparison between the French strains and the OIE-recommended vaccine strains. Amino acid identity to A/equine/Ohio/1/2003 is represented as a dot.

### Comparison of M1, M2, NS1 and NEP proteins

Although eleven substitutions were found (mainly accumulating in M1) between A/equine/Richmond/1/2007 and A/equine/Ohio/1/2003, only ten substitutions were identified between A/equine/Paris/1/2018 and A/equine/Ohio/1/2003 (four in NS1) (**Fig. 7**).

### PB1-F2

The analysis of the gene product PB1-F2, encoded by a +1 reading frame shift of segment 2, showed a large number of substitutions. PB1-F2 is an accessory (nonstructural) protein that presents the highest percentage of substitutions, with twenty-two substitutions for the short versions of PB1-F2 made of 81 amino acids. Interestingly, a stretch of nine residues was present at the C-ter of PB1-F2 encoded by all the predivergent strains [35], but only in a single FC2 virus (A/equine/Saone-et-Loire/1/2015) and in four of the eleven FC1 strains analyzed, suggesting that PB1-F2 functions in equine cells do not need these last amino acid stretches (**Fig. 8)**. While amino acids that have been described to be associated with pathogenicity (T51 and V56; [36]) are conserved among the analyzed strains, residues involved in the inflammatory response (R75 and R79; [37]) are not systematically present. The S66N substitution was identified in all the PB1-F2s analyzed, except those of the predivergent strains, possibly marking a decrease in virulence [38, 39].

**Figure 8:**
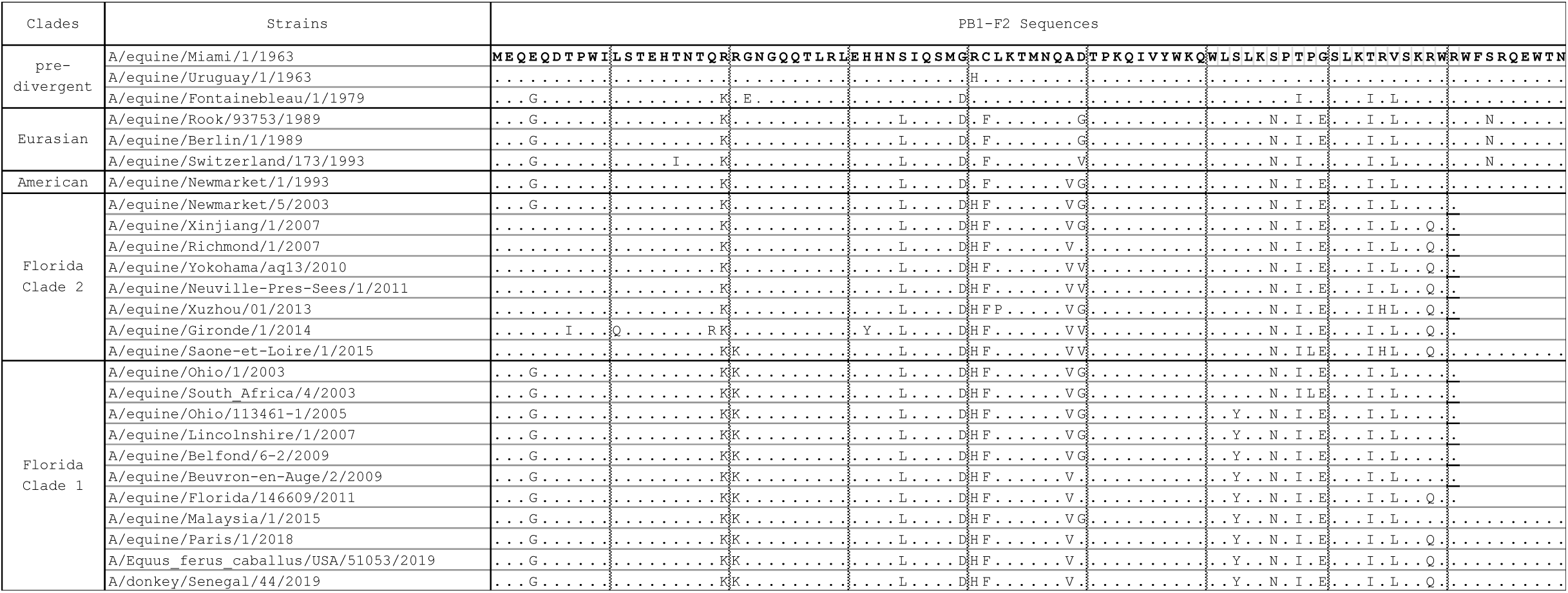
PB1-F2 amino acid sequence comparison. The analysis includes the A/equine/Beuvron-en-Auge/2/2009 and A/equine/Paris/1/2018 strains as well as representative strains. Amino acid identity to A/equine/Miami/1/1963 is represented as a dot.

### Functional characterization of equine PB1-F2

Although PB1-F2 is dispensable for virus replication, it plays significant roles in pathogenesis by altering inflammatory responses, interfering with the host’s innate immune response, and promoting secondary bacterial infections [39–52]. In infected cells, variants of PB1-F2 target mitochondria [42, 46, 53]. Recombinant PB1-F2 has been shown to destabilize and permeabilize synthetic membranes [54–56]. To compare the respective properties of long (90-amino acids long) versus short (81-amino acids long) forms of PB1-F2 of equine viruses, plasmids encoding its A/equine/Paris/1/2018 and the A/equine/Ohio/1/2003 variants were transfected, and their effects on mitochondrial activity were analyzed. **Fig. 9A** shows that the expression of both forms of PB1-F2 resulted in the suppression or lowering of their mitochondrial inner-membrane potential when compared to cells that did not express it, according to a strong decrease in the MitoTracker staining in cells expressing PB1-F2.

**Figure 9:**
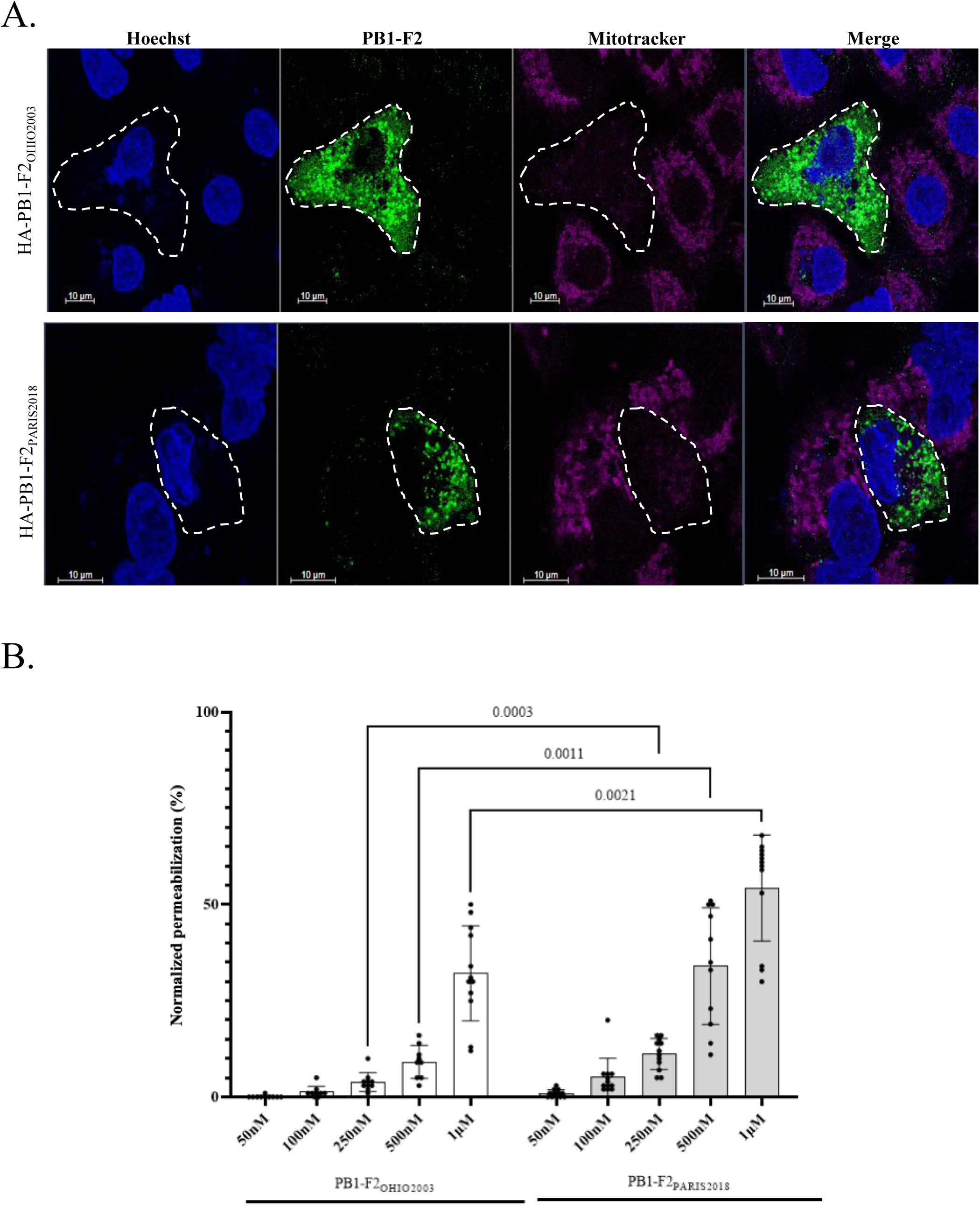
Comparison of biological properties of the virulence factor PB1-F2 of A/equine/Ohio/1/2003 and A/equine/Paris/1/2018. (**A**) Disruption of mitochondrial membrane potential (ΔΨm) in A549 cells expressing HA-tagged PB1-F2 variants from A/equine/Paris/1/2018 (HA-PB1-F2_PARIS2018_) and A/equine/Ohio/1/2003 (HA-PB1-F2_OHIO2003_) viruses. Cells were fixed 48 h post transfection and processed for indirect immunofluorescence staining with an anti-HA-tag rat antibody and an anti-rat secondary antibody coupled with Alexa Fluor 488 (green). Mitochondria were revealed using the ΔΨm-sensitive mitochondrial dye MitoTracker CMX Ros (magenta), and nuclei were revealed with Hoechst (blue). Scale bars, 10 μm. (**B**) Membrane permeabilization assay using recombinant forms of PB1-F2 encoded by A/equine/Paris/1/2018 (PB1-F2_PARIS2018_) and A/equine/Ohio/1/2003 (PB1-F2_OHIO2003_) viruses. LUVs mimicking mitochondrial outer-membrane composition containing the fluorophore probe (ANTS) and quencher (DPX) were incubated with serial dilutions of PB1-F2 forms. The experiment was carried out 4 times in triplicate. Statistical analysis was carried out with REML F(1,99) = 55.01, P<0.0001, and Šídák’s multiple comparison. P values are indicated in the figure.

To further compare the intrinsic properties of the two variants, a lipid vesicle permeabilization assay was used with large unilamellar vesicles (LUVs) composed of synthetic lipid vesicles mimicking the composition of the outer mitochondrial membrane (OMM) [57]. The two PB1-F2 variants were incubated with LUVs containing a fluorescent soluble probe (ANTS) and its quencher (DPX). The permeabilization of LUVs induced ANTS and DPX efflux, which consequently resulted in dilution and dissociation of the fluorescent probe and its quencher in the extravesicular milieu, as revealed by an increase in ANTS fluorescence. **Fig. 9B** shows that both PB1-F2 variants induced permeabilization of the vesicles in a dose-dependent manner, and the specific permeabilization activity of the A/equine/Paris/1/2018 PB1-F2 variant was twofold higher than that of its homolog.

## Discussion

### Whole-genome sequencing

Whole-genome sequences of four equine influenza viruses were produced using a long-read nanopore sequencer on amplicon RT‒PCR products. In addition, these sequences were compared to those obtained by the same sequencer using a direct RNA sequencing library, which could be innovative for characterizing native viral RNA genomes. We confirmed that direct RNA sequencing requires a large amount of RNA material, rendering the accuracy of the sequencing difficult to control [58]. However, when carried out with sufficient material, the two sequencing approaches fulfilled the needed standard for nucleotide sequence determination.

### Phylogeny

The phylogenetic trees of genomic segments confirmed that A/equine/Paris/1/2018 and A/equine/Beuvron-en-Auge/2/2009 belonged to the EIV H3N8 FC1 (**Fig. 3** and)11, 12, 20]) and did not allow the identification of possible segment reassortment events between EIVs. With the integration in our analyses of viruses belonging to FC2, we possibly identified with the phylogenetic tree of segment 7 the mark of a reassortment event in A/equine/Jouars/4/2006 and A/equine/Newmarket/5/2003 between clades 1 and 2 EIV (Fig. 4E). While these two viruses group with FC2 viruses in segments 1-6 and 8, they appear to be more related to FC 1 viruses with their segment 7 sequences. Further investigations are needed to validate this observation.

### Antigenicity

Since 2010, the OIE-ESP has recommended the incorporation of representative EIV strains from both FC1 and FC2 into EI vaccines. Comparison of HA sequences highlights several substitutions between the French EIV strains and the OIE-recommended strain A/equine/Ohio/1/2003 (FC1). The strain A/equine/Paris/1/2018 presents twenty-two substitutions when compared to A/equine/Ohio/1/2003, five of which (A138S, T163I, N188T, R62K, and N63D) are in antigenic sites (site A for the first residue, site B for the two following residues and site E for the last two). The accumulation of these amino acid substitutions within the antibody-binding sites in HA could be sufficient to lead to antigenic drift. We identified one of these substitutions (T163I) only in A/equine/Paris/1/2018 when compared to FC1 and FC2 viruses. According to Wilson and Cox (1990) [59], four or five amino acid substitutions in two separate antigenic sites should be sufficient for escape from preexisting immunity and lead to vaccine failure for human influenza A viruses. For equine influenza A viruses, 10-16 amino acid differences between outbreak and vaccine strains could lead to vaccine breakdown [30, 60]. These results confirm that an EIV clade 1 virus, A/equine/Ohio/1/2003, still constitutes an efficient vaccine strain in recent EIV outbreaks, as previously shown in a large-scale serological study [20]. A similar conclusion could be reached with circulating FC2 EIV and the OIE-recommended vaccine strain A/equine/Richmond/2007, with only 4 substitutions identified in the antigenic sites.

### Equine influenza markers

Equine influenza H3N8 viruses represent a single genetic lineage [61] resulting from the crossover of an avian influenza virus since its first isolation in 1963 [10]. The adaptation of avian influenza A virus to the equine host has been documented, and several host-specific markers have been identified [61, 62]. Comparison between the FluPol, M1, M2, NS1, and NEP sequences of A/equine/Paris/1/2018 and A/equine/Beuvron-en-Auge/2/2009 with representatives of earlier and FC2 strains shows a general conservation of the equine-specific markers with some exceptions. In PB1, a reversion from the recent (since 1997) equine marker I114 was identified in A/equine/Paris/1/2018 (FC1) and A/equine/Saone-et-Loire/1/2015 (FC2) to valine. Additionally, the F94L and K621R substitutions appeared since 2005 in FC1 viruses only. In PA, reversion of the equine E237 to the avian K237 marker has been observed for the most recent Fc1 strain (A/equine/Paris/1/2018) and since 2007 for FC2 strains. This position pertains to a cluster of additional equine-specific markers (positions 213, 216, 217, 231, and 244). S409N substitution was also revealed in A/equine/Paris/1/2018 and A/equine/Beuvron-en-Auge/2/2009, confirming a previously recognized mammal adaptation marker [63] in FC1 viruses [62]. In PB2, the I398V substitution was identified in FC1 viruses in 2005. Similarly, the A684T and A661T substitutions were identified in recent FC1 viruses since 2011 and 2015, respectively. Positions 661 and 684 are known as markers for mammalian adaptation in other influenza viruses [33, 64–67].

### PB1-F2

PB1-F2 is an accessory protein (influenza viruses circulating in humans and other mammalian species do not encode this polypeptide) that is usually 90 amino acids long and displays proinflammatory properties [37]. In mice, amino acids L62, R75, R79, and L82 from influenza A viruses were sufficient to generate an inflammatory response. Mutations at these four positions are sufficient to attenuate the pro-inflammatory properties of the protein. It was thus suggested that some PB1-F2 noninflammatory motifs (P62, H75, Q79, and S82) may diminish the risk of secondary bacterial infection [37]. Moreover, it was experimentally validated that the PB1-F2 proinflammatory motif increased morbidity in primary viral infection and enhanced secondary bacterial infection in mice. Our study as well as [11] shows that the A/equine/Beuvron-en-Auge/2/2009 strain displays a pro-inflammatory motif (L62, R75, and R79) when compared to the A/equine/Paris/1/2018 virus with only L62 and R75. However, another marked difference between these two PB1-F2 is their length. While that of A/equine/Beuvron-en-Auge/2/2009 is only 81 amino acids long, PB1-F2 encoded by A/equine/Paris/1/2018 is 9 amino acids longer with a sequence pattern alternating charged and hydrophobic residues and a hydrophobic residue at position 82, a tryptophan. Full-length versions of PB1-F2 (predominantly 87 or 90 amino acids) have been reported to specifically translocate into mitochondria through their C-terminal region, which acts as a mitochondrial targeting sequence and induces apoptosis [46, 53, 68, 69]. Our functional analyses (on cellular mitochondria and synthetic membranes) reveal a different behavior of the 81- and 90-amino acid-long PB1-F2. Membrane permeabilization was shown to be more efficient with the longer than with the shorter (81 amino acid long) version of PB1-F2 on synthetic membranes. Both forms were able to block the mitochondrial membrane potential when expressed in the cell cytosol. We thus favor the hypothesis that both the length and the amino acid composition may account for the contribution of PB1-F2 in virulence.

## Conclusion

In conclusion, our study highlights the ongoing evolution of equine influenza viruses, with subtle antigenic changes in hemagglutinin and unique genetic variations notably identified in the A/equine/Paris/1/2018 strain. Furthermore, this strain encodes a full-length accessory protein, PB1-F2, resulting in higher permeabilization capacity when compared to shorter forms and possibly contributing to its virulence. The use of advanced long-read sequencing technologies appears to be imperative for monitoring subtle genetic variabilities of emerging variants to identify key virulence markers in the ever-changing landscape of EIV.

## Materials and Methods

### Cell Cultures

A549 cells (human alveolar epithelial cells, American Type Culture Collection) and MDCK cells (Madin-Darby Canine Kidney cells, ATCC) were cultured in minimal essential medium (MEM) (Merck) containing 2 mM L-glutamine, 100 IU/mL penicillin, 100 μg mL−1 streptomycin, and 10% fetal bovine serum. Cells were maintained at 37°C in a 5% CO2 incubator.

### Viruses

Equine influenza viruses (EIV) H3N8 A/equine/Beuvron-en-Auge/2/2009, A/equine/Paris/1/2018, and the vaccine strains A/equine/Richmond/1/2007 and A/equine/South Africa/4/2003 were isolated from sick horses during respiratory disease outbreaks. The nasopharyngeal swabs collected were placed in 5 ml of virus transport medium containing minimum essential medium supplemented with 10% fetal bovine serum and 1% w/v antibiotics (penicillin, streptomycin, and amphotericin).

EIV viruses were first amplified by passaging in 11-day-old embryonated chicken eggs (PA12 White Leghorn strain). Inocula were injected into the allantoid cavity (100 µl per egg). A second virus amplification step was carried out in 25 cm^2^ flasks of MDCK cell monolayers. When cell lysis was observed, cultures were stopped, and RNA extraction was performed immediately.

### RNA extraction

Extraction of EIV RNA from EIV-infected MDCK cells was carried out using TRIzol LS Reagent (Life Technologies) and further purified using the RNeasy MinElute clean-up kit (Qiagen) according to the manufacturer’s recommendations. RNA integrity was assessed on an Agilent 2100 Bioanalyzer using the RNA 6000 nano kit (Agilent, Santa Clara, CA) following the manufacturer’s instructions. We monitored RNA yield and purity with a NanoDrop ND-2000c spectrophotometer.

### MinION long-read library preparation, sequencing and data analysis

#### cDNA synthesis

Purified RNA was reverse transcribed using SuperScript III (Thermo Scientific) and primers designed by Keller et coll. and complementary to the conserved 3’ end of influenza A vRNA [33]. We used primers RTA-U12 (5’-AGCAAAAGCAGG) expected to target the segments PA, NP, M, NS and RTA-U12.4 (5’-AGCGAAAGCAGG) expected to target the segments PB2, PB1, HA, NA, combined in a 2:3 molar ratio [33]. 500 ng of total RNA and 10 pmol of specific primers (2:3 molar ratio RTA-U12, RTA-12.4) were denatured for 5 min at 65°C, centrifuged, and stored on ice before adding the reaction mix, according to the manufacturer’s instructions. We incubated the RT reactions at 25°C for 10 min and then 50°C for 60 min. The reaction was then stopped by heating at 70°C for 15 min. After cDNA synthesis, RNA was degraded by incubation with 2 U of RNase H for 20 min at 37°C. The RNA hydrolysis reaction was stopped by heating at 70°C for 10 min, and the cDNAs were stored at –20°C until use. We evaluated the quantity and quality of cDNA on sixfold dilutions with the RNA 6000 Pico kit (Agilent) on an Agilent 2100 Bioanalyzer.

#### cDNA amplification

The eight influenza A genomic segments were amplified by PCR using the cDNA previously produced. Platinum II Taq Hot start DNA-polymerase (Invitrogen) was used according to the manufacturer’s instructions, with primers set complementary to the 5’ and 3’ ends of each influenza A genome segment (**Supp. Table 3**). Amplified DNA products were purified using AMPure XP beads (Beckman Coulter Inc., Pasadena, CA, USA) at a ratio of 1.2:1 volume of beads per sample, and DNA yield was monitored with a NanoDrop ND-2000c spectrophotometer and a Qubit fluorimeter using a Qubit dsDNA BR kit (Invitrogen).

#### Nanopore sequencing and data analysis

For each of the four strains, the eight purified PCR products were pooled at an equimolar ratio and used as input for library generation using the Ligation Sequencing Kit SQK-LSK109 and the Native Barcoding Expansion 1-12 kit EXP-NBD104 according to the manufacturer’s instructions (Oxford Nanopore Technologies). The barcode-ligated DNA samples were pooled at an equimolar ratio and used for final adapter ligation. We loaded 50 fmol of the purified adapter-ligated DNA library onto a MinION Flow-cell (R9.4.1; FLO-MIN106D) and run it on a MinION Mk1C device according to the manufacturer’s instructions. Guppy (version 5.1.13) was used for basecalling and demultiplexing. Nanofilt (version 2.8.0) was used to filter reads based on their size and quality: size > 600 bp, Q>10. The filtered reads were mapped using minimap2 (version 2.22) and A/equine/Ohio/113461-1/2005 as the reference genome (GenBank accession numbers: CY067323, CY067324, CY067325, CY067326, CY067327, CY067328, CY067329, CY067330). SAMtools (version 1.14) was used to convert the data into bam and medaka (version 1.4.4) for variant calling. Finally, the Integrative Genomics viewer desktop application (IGV, version 2.16.2) was used for visualization.

The newly sequenced viral genomes have been deposited in the European Nucleotide Archive under project accession number X, available at www.ebi.ac.uk/ena.

### Sequence multialignment and phylogenetic trees

A multiple alignment of all nucleotide sequences of the eight genes of equine influenza of type A H3N8 was obtained using the Muscle algorithm and the maximum likelihood method and Hasegawa-Kishino-Yano model [70]. The tree with the highest likelihood is shown. The percentage of replicate trees in which the associated taxa clustered together in the bootstrap test 1000 replicates [71] are shown next to the branches. Initial tree(s) for the heuristic search were obtained automatically by applying neighbor-joining and BioNJ algorithms to a matrix of pairwise distances estimated using the maximum composite likelihood (MCL) approach and then selecting the topology with superior log likelihood value. A discrete Gamma distribution was used to model evolutionary rate differences among sites (5 categories (+G, parameter)). The codon positions included were 1st+2nd+3rd+Noncoding. Evolutionary analyses were conducted in MEGA11 [72, 73]. All accession numbers are listed in **Supp. Table 1**.

The amino acid sequences of viral proteins (PB2, PB1, PB1-F2, PA, PA-X, HA, NP, NA, M1, M2, NS1, and NEP) from recent EIV isolated in France were aligned with the strain A/equine/Ohio/1/2005 used as a reference for consensus sequence construction using Clustal Omega from EMBL-EBI [74] and Unipro UGENE [75].

### Plasmids

Codon-optimized open reading frames encoding HA-tagged versions of PB1-F2 of viral strains A/equine/Ohio/1/2003 and A/equine/Paris/1/2018 were cloned in the eukaryotic expression vector pCAGGS at the Not I and Bgl II restriction sites. Codon-optimized open reading frames encoding His-tagged versions of PB1-F2 of A/equine/Ohio/1/2003 and A/equine/Paris/1/2018 were cloned in the bacterial expression vector pET-28a+ at the Nde I and Xho I restriction sites.

### Immunohistochemistry - Confocal Microscopy

A549 cells were seeded at 0.5 x 10^6^ cells per well on 18 mm diameter glass lamellas and incubated for 24 h at 37°C and 5% CO2. Cells at 80∼90% confluence were transfected with 200 ng of pCAGGS derivates using Lipofectamine® 2000 (11668027, Thermo Fisher Scientific) following the manufacturer’s instructions. Forty hours after transfection, MitoTracker CMX Ros (M7512, Thermo Fisher Scientific) was added to the cell culture at a final concentration of 500 nM for 30 min. Next, after cell culture medium removal, the cells were fixed using 4% paraformaldehyde for 30 min at room temperature (RT). Cell monolayers were washed in phosphate saline buffer (PBS) and PBS completed with 0.1% Triton X-100 (PBS-Tx) and with 1% w/v bovine serum albumin (BSA) for 1 h at RT. The cells were then incubated with a rabbit anti-HA-tag antibody (H6908, Sigma‒Aldrich) in PBS-Tx supplemented with 0.2% BSA. After three washes in PBS-Tx, an anti-rabbit immunoglobulin goat antibody labeled with Alexa Fluor 488 (A11008, Invitrogen, OR, USA) in PBS-Tx completed with 0.2% BSA was added for 2 h at RT. Nuclei were marked with Hoechst diluted to 1/100 in PBS 1x for 5 min at RT. Subcellular localization images were taken using a Zeiss LSM 700 confocal 187 microscope with a x63 objective.

### PB1-F2 production in *E. coli* and purification

BL-21 Rosetta cells (Stratagene) were transformed with the resulting plasmids and cultured to an optical density (OD) of 0.8 before overnight incubation at 28°C in 1 mM isopropyl 1-thio-β-D-galactopyranoside (IPTG) under agitation. Next, bacteria were pelleted and resuspended in 50 mM Tris (pH 7.4), 10 mM EDTA, and 0.1% Triton X-100 buffer and incubated at 37°C for 30 min. The suspension was sonicated and centrifuged at 10.000 × g for 30 min at 4°C. Pellets were resuspended in solubilization buffer (20 mM Tris (pH 7.4), 0.5 M NaCl, 5 mM imidazole, and 8 M urea) and centrifuged at 10,000 × g for 30 min at 4°C. Supernatants were sonicated and filtrated using 0.8 µm filters (SLAAR33S, MilliporeSigma™ Millex™) before loading on a Histrap FF IMAC column (17531901, Cytiva) using the AKTA Purifier 100 FPLC chromatographic system (GE Healthcare). Fractions containing PB1-F2 were pooled and subjected to size exclusion chromatography on a Sepharose S200 column equilibrated with solubilization buffer. Next, urea was removed from the S200 PB1-F2-containing fractions on a 53 mL HiPrep™ 26/10 Sephadex G-25 resin column (GE17-5087-01, Sigma‒Aldrich) equilibrated with 5 mM ammonium acetate buffer, pH 5. Fractions containing PB1-F2 were lyophilized and stored at −20°C. Prior to their use, lyophilized PB1-F2 powder was dissolved in 5 mM sodium acetate buffer (pH 5). Protein concentration was estimated by measuring OD at 280 nm and using the extinction coefficients of 23490 M^−1^cm^−1^ for the Ohio protein and 37470 M^−1^cm^−1^ for its Paris homolog.

### Lipid vesicle preparation

(16:0-18:1) 1-Palmitoyl-2-oleoyl-sn-glycero-3-phospho-L-serine (POPS) (840034), 1-palmitoyl-2-oleoyl-sn-glycero-3-phospho-(1’-rac-glycerol) (PG) (840457), and (18:1) cardiolipin 1’,3’-bis[1,2-dioleoyl-sn-glycero-3-phospho]-glycerol (DOCL) (840044) were purchased from Avanti Polar Lipids (Alabaster, AL, USA). (16:0-18:1) 1-Palmitoyl-2-oleoyl-glycero-3-phosphocholine (POPC) (37-1618-9), (16:0-18:1) 1-palmitoyl-2-oleoyl-sn-glycero-3-phosphoethanolamine (POPE) (37-1828-7), and soybean L-α-phosphatidylinositol (PI) (37-0130-7) were purchased from Larodan (France). ANTS (FP-46574B, 8-aminonapthalene-1,3,6 trisulfonic acid) and DPX (FP-47017A, p-xylene-bis-pyridinium bromide) were purchased from Interchim (Montluçon, France). Sodium acetate buffers and phosphate buffers were of analytical grade. Reagents for SDS‒PAGE electrophoresis were obtained from Invitrogen (France).

Lipids POPC, POPE, POPS, PI, and DOCL were used at a molar ratio of 5.5:2.5:1.5:1:0.5 to mimic mitochondria outer membranes (OMM). The mix of lipids with 20 mM ANTS (fluorophore probe) and 60 mM DPX (quencher) in a final concentration of 10 mM sodium acetate (pH 5) was sonicated using a sonicator tip to obtain an emulsion. Reversed-phase evaporation was carried out using a Heidolph Laborota 4003 apparatus to obtain large unilamellar vesicles (LUVs). LUV preparations were extruded three times through a Swinny filter (XX3001200, Millipore) using polycarbonate filters with pore size diameters of 1.2 μm, 0.4 μm and 0.2 μm (Merck Millipore, Darmstadt, Germany). Unencapsulated ANTS and DPX were removed by gel filtration through a 5 mL HiTrap Desalting Sephadex G-25 resin column (GE Healthcare Life Sciences). To ensure the correct size and obtain LUVs, dynamic light scattering (DLS) measurements were performed on a Nano series Zetasizer (Malvern Instruments, Paris, France).

### Lipid vesicle permeabilization assay

For permeabilization assays, LUVs were incubated at 0.4 mM lipid concentration in 10 mM sodium acetate (pH 5) at 25°C in a black p96-well plaque (Greiner), and fluorescence titrations were performed with an FP-8200 Jasco spectrofluorometer equipped with a Peltier-thermostated ETC-272T (25°C). The excitation wavelength was set at 360 nm, and the emission of ANTS was measured between 500-600 nm at a bandwidth of 5 nm to ensure that the signal perceived was indeed permeabilization and not unspecific diffraction. The intensity was measured before and after the addition of PB1-F2 at final concentrations of 1 μM, 500 nM, 250 nM, 100 nM, and 50 nM. The maximum intensity of permeabilization, corresponding to the maximum intensity of ANTS fluorescence, was measured after the addition of 0.1% (v/v) Triton X-100. The experiment was carried out 4 times in triplicate. Statistical analysis was carried out with REML F(1,99) = 55.01, P<0.0001 and Šídák’s multiple comparison (1 μM P value = 0.0021; 500 nM P value = 0.0011; 250 nM P value=0.0003) on Prism v9.

## Declarations

## Availability of data and materials

The datasets used and/or analyzed during the current study are available from the corresponding author upon reasonable request.

## Acknowledgments

We thank Christophe Chevalier for discussion and advice. We also thank the MIMA2 platform (https://doi.org/10.15454/1.5572348210007727E12) for access to confocal microscopy.

## Author contributions

Conceptualization: Loïc Legrand, Eric Barrey, Stéphane Pronost, Bernard Delmas

Investigation: Lena Kleij, Elise Bruder, Dorothée Raoux-Barbot, Nathalie Lejal, Quentin Nevers, Bruno Da Costa, Charlotte Deloizy, Loïc Legrand, Eric Barrey, Alexandre Chenal, Stéphane Pronost, Bernard Delmas, Sophie Dhorne-Pollet

Methodology: Lena Kleij, Elise Bruder, Dorothée Raoux-Barbot, Nathalie Lejal, Quentin Nevers, Bruno Da Costa, Charlotte Deloizy, Sophie Dhorne-Pollet

Formal analysis: Lena Kleij, Quentin Nevers, Bruno Da Costa, Charlotte Deloizy, Loïc Legrand, Eric Barrey, Alexandre Chenal, Stéphane Pronost, Bernard Delmas, Sophie Dhorne-Pollet

Funding acquisition: Loïc Legrand, Eric Barrey, Bernard Delmas

Supervision: Loïc Legrand, Eric Barrey, Alexandre Chenal, Stéphane Pronost, Bernard Delmas, Sophie Dhorne-Pollet

Visualization: Lena Kleij, Quentin Nevers, Sophie Dhorne-Pollet Writing – original draft: Lena Kleij, Bernard Delmas

Writing – review & editing: Lena Kleij, Quentin Nevers, Eric Barrey, Alexandre Chenal, Stéphane Pronost, Sophie Dhorne-Pollet, Bernard Delmas

