## Supplementary information for "Genome sequences of equine influenza A subtype H3N8 viruses by long read sequencing and functional characterization of the PB1-F2 virulence factor of A/equine/Paris/1/2018"

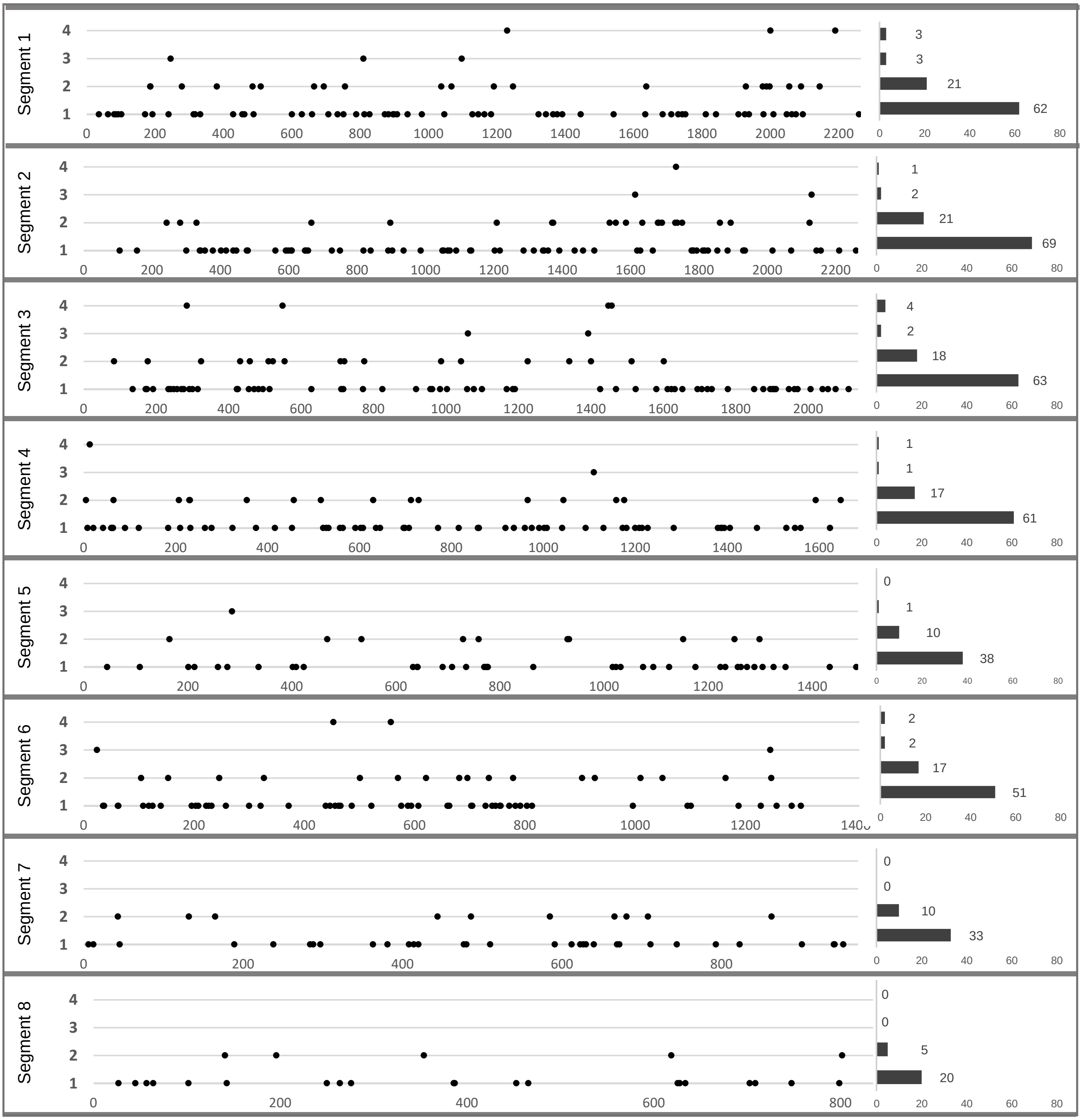


Supp Fig 1 KLEIJ et al., 2023

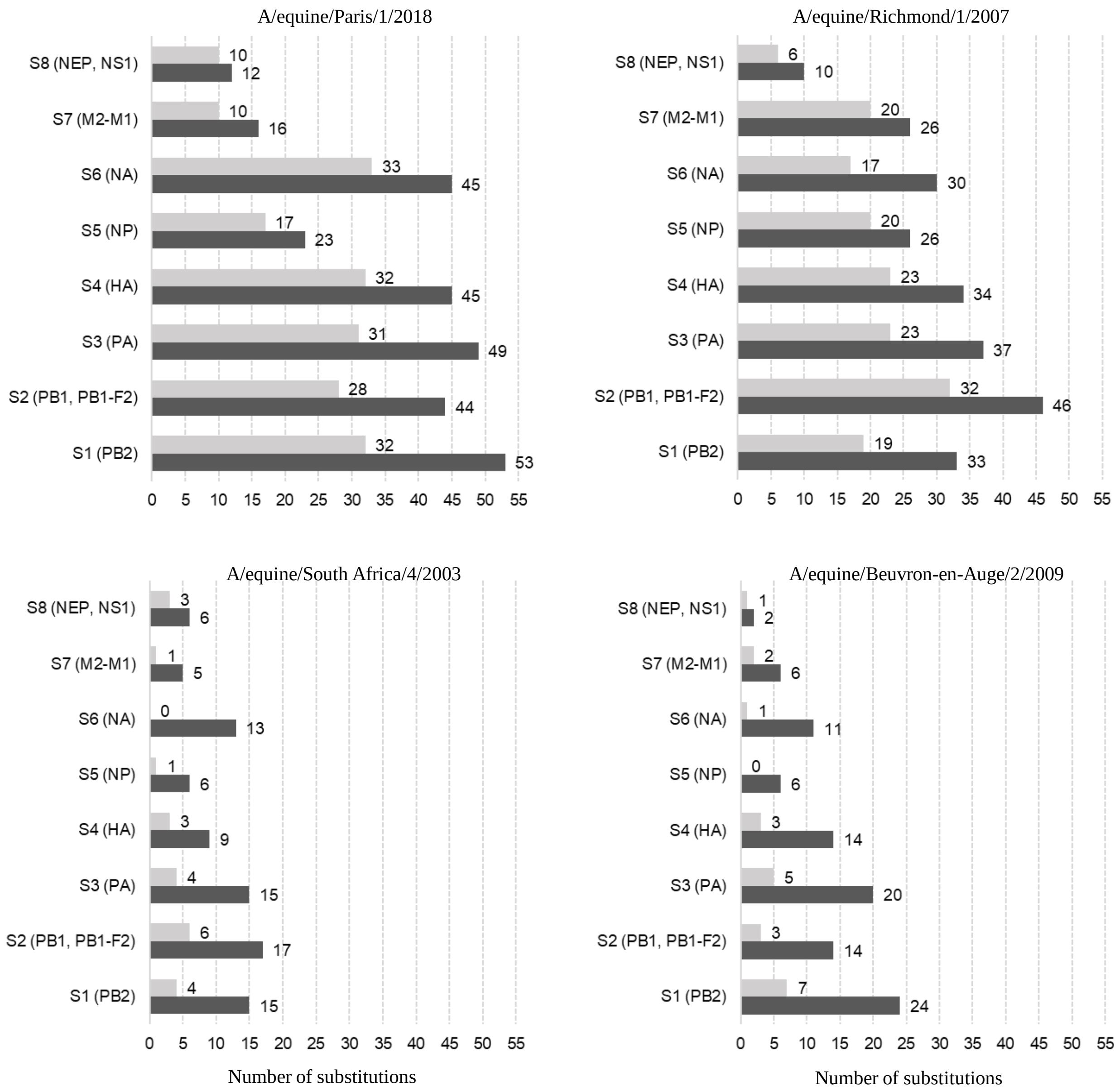


**Supp Fig 2 KLEIJ et al., 2023**

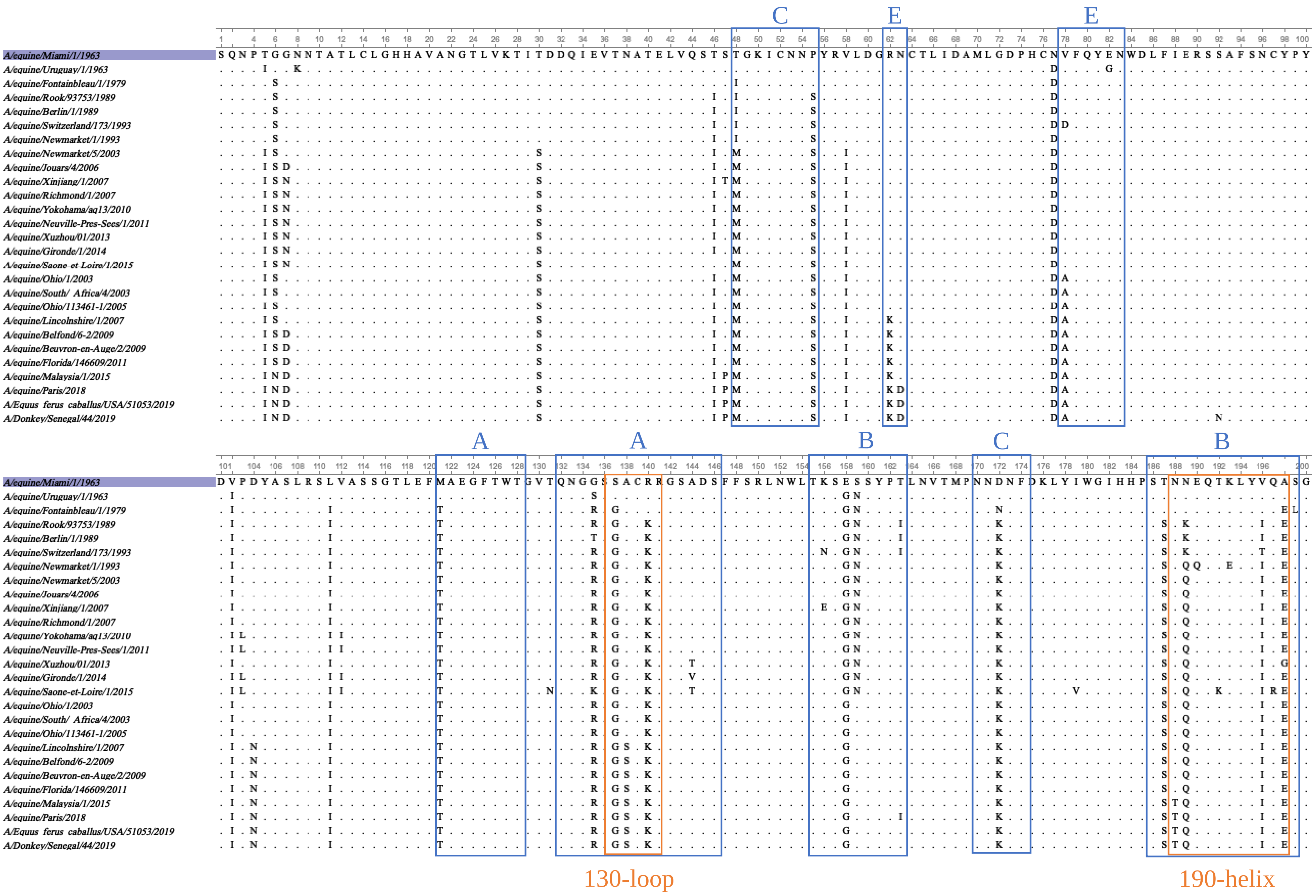


**Supp Fig 3 KLEIJ et al., 2023**

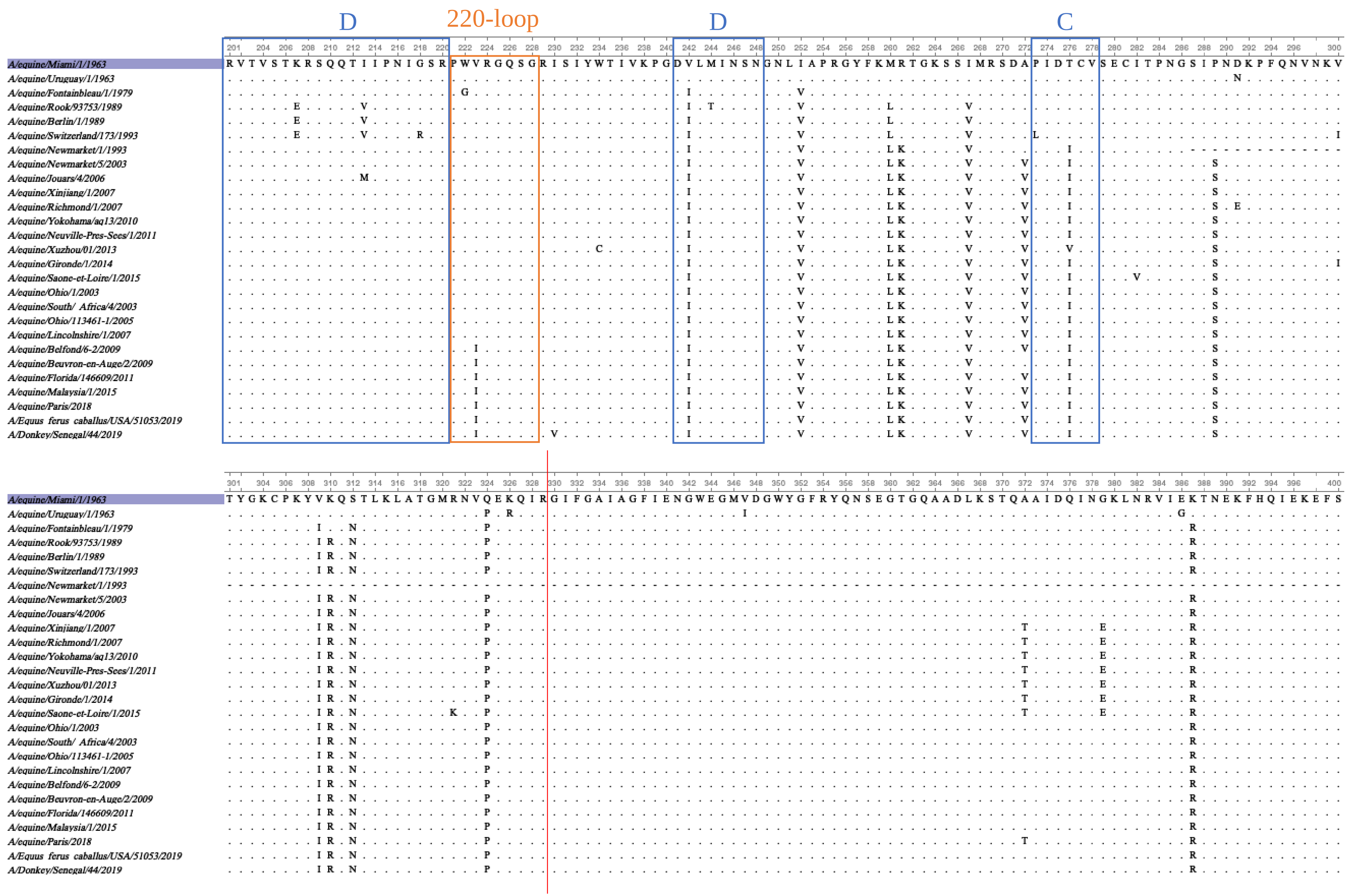


Supp Fig 3 KLEIJ et al., 2023

```
                                     401                                                                                                500
A/equine/Miami/1/1963                EVEGRIQDLEKYVEDTKIDLWSYNAELLVALENQHTIDLTDAEMNKLFEKTRRQLRENAEDMGNGCFKIYHKCDNACIESIRNGTYDHDIYRDEALNNRF
A/equine/Uruguay/1/1963              ...........................................................................................K........
A/equine/Fontainbleau/1/1979         ...............................................................G..............G.........Y...........
A/equine/Rook/93753/1989             ...............................................................G..............G.........Y...........
A/equine/Berlin/1/1989               ...............................................................G..............G.........Y...........
A/equine/Switzerland/173/1993        ...............................................................G..............G.........Y...........
A/equine/Newmarket/1/1993            ----------------------------------------------------------------------------------------------------
A/equine/Newmarket/5/2003            ...............................................................G..............G.........Y...........
A/equine/Jouars/4/2006               ....K....................G.....................................G..............G.........Y...........
A/equine/Xinjiang/1/2007             ...............................................................G..............G.........Y...........
A/equine/Richmond/1/2007             ...............................................................G..............G.........Y...........
A/equine/Yokohama/aq13/2010          ...............................................................G..............G.........Y...........
A/equine/Neuville-Pres-Sees/1/2011   ...............................................................G..............G.........Y...........
A/equine/Xuzhou/01/2013              ...............................................................G..............G.........Y...........
A/equine/Gironde/1/2014              ...............................................................G..............G.........Y...........
A/equine/Saone-et-Loire/1/2015       ...............................................................G..............G.........Y...........
A/equine/Ohio/1/2003                 ...............................................................G..............G.........Y...........
A/equine/South/ Africa/4/2003        ...............................................................G..............G.........Y.....X.....
A/equine/Ohio/113461-1/2005          ...............................................................G..............G.........Y...........
A/equine/Lincolnshire/1/2007         ...............................................................G..............G.........Y...........
A/equine/Belfond/6-2/2009            ...............................................................G..............G.........Y...........
A/equine/Beuvron-en-Auge/2/2009      ...............................................................G..............G.........Y...........
A/equine/Florida/146609/2011         ...............................................................G..............G.........Y...........
A/equine/Malaysia/1/2015             ...............................................................G.........N....G.........Y...........
A/equine/Paris/2018                  .................................................R...L.........G.........N....G.........Y...........
A/Equus ferus caballus/USA/51053/2019 ...............................................R...L.........G.........N....G.........Y...........
A/Donkey/Senegal/44/2019             .................................................R...L.........G.........N....G.........Y...........
```

```
                                     501                                              550
A/equine/Miami/1/1963                QIRGVELKSGYKDWILWISFAISCFLICVVLLGFIMWACQKGNIRCNICI
A/equine/Uruguay/1/1963              ..................................................
A/equine/Fontainbleau/1/1979         ..K...............................................
A/equine/Rook/93753/1989             ..K...............................................
A/equine/Berlin/1/1989               ..K...............................................
A/equine/Switzerland/173/1993        ..KS..............................................
A/equine/Newmarket/1/1993            --------------------------------------------------
A/equine/Newmarket/5/2003            ..K...............................................
A/equine/Jouars/4/2006               ..K...............................................
A/equine/Xinjiang/1/2007             ..K...............................................
A/equine/Richmond/1/2007             ..K...............................................
A/equine/Yokohama/aq13/2010          ..K...............................................
A/equine/Neuville-Pres-Sees/1/2011   ..K...............................................
A/equine/Xuzhou/01/2013              ..K...............................................
A/equine/Gironde/1/2014              ..K.......N....M.......N................----------
A/equine/Saone-et-Loire/1/2015       ..K.......................V.......................
A/equine/Ohio/1/2003                 ..K...............................................
A/equine/South/ Africa/4/2003        ..K...............................................
A/equine/Ohio/113461-1/2005          ..K...............................................
A/equine/Lincolnshire/1/2007         ..K.......................V.......................
A/equine/Belfond/6-2/2009            ..K.......................V.......................
A/equine/Beuvron-en-Auge/2/2009      ..K.......................V.......................
A/equine/Florida/146609/2011         ..K.......................V.......................
A/equine/Malaysia/1/2015             ..K...............................................
A/equine/Paris/2018                  ..K...............................................
A/Equus ferus caballus/USA/51053/2019 ..K...............................................
A/Donkey/Senegal/44/2019             ..K...............................................
```

**Supp Fig 3 KLEIJ et al., 2023**

```
                                              1-10       11-20      21-30      31-40      41-50      51-60      61-70      71-80      81-90      91-100
A/equine/Miami/1/1963                         MNPNQKIITI GSASLGLVIL NVILHVVSII VTVLVLSNNG TGPNCNGTII REYNETVRVE RITQWYNTNI IEYIEEPSNE YYMSNTEPLC EAQGFAPFSK
A/equine/Uruguay/1/1963                       .......... ........F. .......... .......... .......... ........I. .......... .....R.... .......... ..........
A/equine/Fontainbleau/1/1979                  ........A. ......IL.. .......... ......N... ..L....... .......... .........T .....R.... ...N...... ..........
A/equine/Rook/93753/1989                      ........A. ......IL.. .......... ......N... ..L....... .......... ........ST .....R.... ...N...... ..........
A/equine/Berlin/1/1989                        ........A. ......IL.. .......... ......N... ..L....... .......... ........ST .....R.... ...N...... ..........
A/equine/Switzerland/173/1993                 ........A. ......VL.. .......... ......N... ..L....... .......... ........ST .....RS... ...N...... ..........
A/equine/Newmarket/1/1993                     ........A. ......IL.I .......... ......N... ..L..K.... .......... ........ST .....R.... ...N...... ..........
A/equine/Newmarket/5/2003                     ........A. .F....IL.I .......... ......N... ..L..K.... .......... K.......SA .K...R.P.. ...N...... ..........
A/equine/Jouars/4/2006                        ---------- ----..IL.I .......... ......N... ..L..K.... .......... K....H..SA .K...R.P.. ...N...... ..........
A/equine/Xinjiang/1/2007                      .......... ......IL.I .......... ......N..E ..L..K.... .......... K....H..SA .K...R.P.. ...N...... ..........
A/equine/Richmond/1/2007                      .......... ......IL.I .......... ......N..E ..L..K.... .......... K....H..SA .K...R.P.. ...N...... ..........
A/equine/Yokohama/aq13/2010                   .......... ......IL.I .......... ......N..E ..L..K.... .......... K....H..SA .K...R.P.. ...N...... ..........
A/equine/Neuville-Pres-Sees/1/2011            .......... ......IL.I .......... ......N..E ..L..K.... .......... K....H..SA .K...R.P.. ...N...... ..........
A/equine/Xuzhou/01/2013                       ......K... ......IL.I .........V ......N..E ..L..R.... .......... K....H..SA .K...R.P.. ...N...... ..........
A/equine/Gironde/1/2014                       .......... ......IL.I ....N..... ......N..E .CL..K.M.. .......... K....H..SA .K...R.P.. ...N...... ..........
A/equine/Saone-et-Loire/1/2015                .......... ......IL.I .......... ......N..E ..L..K.... .......... K....H..SA .K...R.P.. ...N...... ..........
A/equine/Ohio/1/2003                          ........A. .F....IL.I .......... ......N..R .DL..K.... .......... K.......ST .K...R.... ...N...... ..........
A/equine/South Africa/4/2003                  ........A. .F....IL.I .......... ......N..R .DL..K.... .......... K.......ST .K...R.... ...N...... ..........
A/equine/Ohio/113461-1/2005                   .......MA. .F....IL.I .......... .....AN..R .DL..K.... .......... K.......ST .K...R.... ...N...... ..........
A/equine/Lincolnshire/1/2007                  ........A. .F....IL.I .......... .....AN..K .DL..K.... .......... K.......SA .K...R.... ...N...... ..........
A/equine/Beuvron-en-Auge/2/2009               .......MA. .F....IL.I .......... .....AN..R .DL..K.... .......... K.......ST .K...R.... ...N...... ..........
A/equine/Belfond/6-2/2009                     .......MA. .F....IL.I .......... .....AN..R .DL..K.... .......... K.......ST .K...R.... ...N...... ..........
A/equine/Florida/146609/2011                  .......MA. .F....IL.I .......... .....AN..R .DL..K.... .......... K.......ST .K...R.... ...N...... ..........
A/equine/Malaysia/1/2015                      ........A. .F....IL.I .......... .....AN..R .DL..K.... .......... K.......ST .K...R.... ...N...... ..........
A/equine/Paris/1/2018                         ........A. .FT...IL.I S......... .....AN..R .DL..KE... .......... K......IST .K.M.K.... ...N...... ..........
A/Equus ferus caballus/USA/51053/2019         ........A. .FT...IL.I S......... .....AN..R .DL..KE... .......... K......IST .K..DK.... ...N...... ..........
A/donkey/Nigeria/VRD19/ZM20 T10 19RS459-5/2019 ........A. .FT...IL.I S......... .....AN..R .DL..KE... .......... K......IST .K...K.... ...N...... ..........

                                              101-110    111-120    121-130    131-140    141-150    151-160    161-170    171-180    181-190    191-200
A/equine/Miami/1/1963                         DNGIRIGSRG HVFVIREPFV SCSPLECRTF FLTQGSLLND KHSNGTVKDR SPYRTLMSVE VGQSPNVYQA RFEAVAWSAT ACHDGKKWMT VGVTGPDAQA
A/equine/Uruguay/1/1963                       .......... .......... .......... .......... .......... .......... .......... .......... .......... ..........
A/equine/Fontainbleau/1/1979                  .......... .......... .......... ...H...... .......... .........K .......... .......... .......... .......T..
A/equine/Rook/93753/1989                      .......... .......... ....S..... .......... .......... .........K I......... ...S...... .......... .......N..
A/equine/Berlin/1/1989                        .......... .......... ....S..... .......... .......... .........K I......... ...S...... .......... .......N..
A/equine/Switzerland/173/1993                 .......... .......... ....S..... .......... .......... .........K I......... ...S...... .......... .......N..
A/equine/Newmarket/1/1993                     .......... .......... ....S..... .......... .......... .........K I......... ...S...... .......... .......N..
A/equine/Newmarket/5/2003                     .......... .......... ....S..... .......... .......... .........K I......... ...S...... .......... .......N..
A/equine/Jouars/4/2006                        .......... .......... ....S..... .......... .......... .........K I......... ...S...... .......... I......N..
A/equine/Xinjiang/1/2007                      .......... .......... ....S..... .......... .......... .........K I......... ...S...... .......... I......N..
A/equine/Richmond/1/2007                      .......... .......... ....S..... .......... .......... .........K I......... ...S...... .......... I......N..
A/equine/Yokohama/aq13/2010                   ........K. .......... ....S..... .......... .......... .........K I......... ...S...... .......... I......N..
A/equine/Neuville-Pres-Sees/1/2011            ........K. .......... ....S..... .......... .......... .........K I......... ...S...... .......... I......N..
A/equine/Xuzhou/01/2013                       .......... .......... ....S..... .......... .......... .........K I......... ...S...... .......... I......N..
A/equine/Gironde/1/2014                       ........K. .......... ....S..... .......... .......... .........K I......... ...S...... .......... I......N..
A/equine/Saone-et-Loire/1/2015                ........K. .......... ....S..... .......... .......... .........K I......... ...S...... .......... I......N..
A/equine/Ohio/1/2003                          .......... .......... ....S..... .......... .......... .........K I......... ...S...... .......... .......N..
A/equine/South Africa/4/2003                  .......... .......... ....S..... .......... .......... .........K I......... ...S...... .......... .......N..
A/equine/Ohio/113461-1/2005                   .......... .......... ....S..... .......... .......... .........K I......... ...S...... .....R.... .......N..
A/equine/Lincolnshire/1/2007                  .......... .......... ....S..... .......... .......... .........K I......... ...S...... .......... .......N..
A/equine/Beuvron-en-Auge/2/2009               .......... .......... ....S..... .......... .......... .........K I......... ...S...... .......... .......N..
A/equine/Belfond/6-2/2009                     .......... .......... ....S..... .......... .......... .........K I......... ...S...... .......... .......N..
A/equine/Florida/146609/2011                  .......... .......... ....S..... .......... .......... .........K I......... ...S...... .......... .......N..
A/equine/Malaysia/1/2015                      .......... .......... ....S..... .......... .......... .........K I......... ...S...... .......... .......N..
A/equine/Paris/1/2018                         .......... .......... ....S..... .......... ......I... .........K I......... ...S...... .......... .......N..
A/Equus ferus caballus/USA/51053/2019         .......... .......... ....S..... .......... ......I... .........K I......... ...S...... .......... .......N..
A/donkey/Nigeria/VRD19/ZM20 T10 19RS459-5/2019 .......... .......... ....S..... .......... ......I... .........K I......... ...S...... .......... .......N..
```

Supp Fig 4 KLEIJ et al., 2023

```
                                          201                                                                                               300
A/equine/Miami/1/1963                     VAVVHYGGVPVDVINSWAGDILRTQESSCTCIKGDCYWVMTDGPANRQAQYRIFKAKDGRIIGQTDINFNGGHIEECSCYPNEGKVECVCRDNWTGTNRP
A/equine/Uruguay/1/1963                   ..................................................................V.................................
A/equine/Fontainbleau/1/1979              ............I....................................K..................................................
A/equine/Rook/93753/1989                  ....N.......I....................................K.................S********************I...........
```

```
                                          201                                                                                               300
A/equine/Miami/1/1963                     VAVVHYGGVPVDVINSWAGDILRTQESSCTCIKGDCYWVMTDGPANRQAQYRIFKAKDGRIIGQTDINFNGGHIEECSCYPNEGKVECVCRDNWTGTNRP
A/equine/Uruguay/1/1963                   ..................................................................V.................................
A/equine/Fontainbleau/1/1979              ............I....................................K..................................................
A/equine/Rook/93753/1989                  ....N.......I....................................K.................S....................I...........
A/equine/Berlin/1/1989                    ....N.......I....................................K.................S....................I...........
A/equine/Switzerland/173/1993             ....N.......I....................................K.................S....................I...........
A/equine/Newmarket/1/1993                 I...N.......I....................................K.................S....................I...........
A/equine/Newmarket/5/2003                 I...N.......I.....................N..............K..........V......S....................I...........
A/equine/Jouars/4/2006                    I...N.......I.....................N..............K..........V......S....................I...........
A/equine/Xinjiang/1/2007                  I...N...I...I....E................N..............K..........V......S....................I...........
A/equine/Richmond/1/2007                  I...N.......I.....................N..............K..........V......S....................I...........
A/equine/Yokohama/aq13/2010               I...N.......I.....................N..............K..........V......S....................I...........
A/equine/Neuville-Pres-Sees/1/2011        I...N.......I.....................N..............K..........V......S....................I...........
A/equine/Xuzhou/01/2013                   T...N...I...I....E................N..............K..........V......S....................I...........
A/equine/Gironde/1/2014                   I...N.......I.....................N..............K..........V......S....................I...........
A/equine/Saone-et-Loire/1/2015            I...N.......I.....................N..............K..........V......S....................I...........
A/equine/Ohio/1/2003                      I...N.......I....................................K..........V......S..E.................I...........
A/equine/South  Africa/4/2003             I...N.......I....................................K..........V......S....................I...........
A/equine/Ohio/113461-1/2005               I...N.......I....................................K..........V......S....................I...........
A/equine/Lincolnshire/1/2007              I...N.......I....................................K..........V......S....................I...........
A/equine/Beuvron-en-Auge/2/2009           I.I.N.......I....................................K.........KV......S....................I...........
A/equine/Belfond/6-2/2009                 I.I.N.......I....................................K.........KV......S....................I...........
A/equine/Florida/146609/2011              I...N.......I....................................K.........KV......S....................I...........
A/equine/Malaysia/1/2015                  I...N.......I....................................K.K.....N.KV....N.S....................I...........
A/equine/Paris/1/2018                     I...N.......I....................................K.K.....N.KV......S....................I...........
A/Equus  ferus caballus/USA/51053/2019    I...N.......I....................................K.K.....N.KV......S....................I...........
A/donkey/Nigeria/VRD19/ZM20 T10 19RS459-5/2019 I...N.....I.I....................................K.K.....N.KV......S....................I...........
```

```
                                          301                                                                                               400
A/equine/Miami/1/1963                     VLVISPDLSYTVGYLCAGIPTDTPRGEDSQFTGSCTSPLGSQGYGVKGFGFRQGNDVWAGRTISRTSRSGFEIIKIRNGWTQNSKDQIRKQVIVDNLNWS
A/equine/Uruguay/1/1963                   ....................................................................................................
A/equine/Fontainbleau/1/1979              I.......................................N.......................................I...................
A/equine/Rook/93753/1989                  I....S..................................NK............T........G.........................R...I......
A/equine/Berlin/1/1989                    I....S..................................NK............T..................................R...I......
A/equine/Switzerland/173/1993             I....S..................................NK............T..................................R...I......
A/equine/Newmarket/1/1993                 I....S..................................NK............T..................................R...I..P...
A/equine/Newmarket/5/2003                 I....S..................................NK............T..................................R...I.DP...
A/equine/Jouars/4/2006                    I....S..................................NK............T..................................R...I.DP...
A/equine/Xinjiang/1/2007                  I....S..................................NK............T..................................R...I.DP...
A/equine/Richmond/1/2007                  I....S..................................NK............T..................................R...I.DP...
A/equine/Yokohama/aq13/2010               I....S..................................NK............T..................................R...I.DP...
A/equine/Neuville-Pres-Sees/1/2011        I....S..................................NK............T..................................R...I.DP...
A/equine/Xuzhou/01/2013                   I....S..................................NK............T..................................R...I.DQ...
A/equine/Gironde/1/2014                   I....S..................................NK............T..................................R...I.DP...
A/equine/Saone-et-Loire/1/2015            I....S..................................NK............T.........................I........R...I.DP...
A/equine/Ohio/1/2003                      I....S..................................NK............T..................................R...I.DP...
A/equine/South  Africa/4/2003             I....S..................................NK............T..................................R...I.DP...
A/equine/Ohio/113461-1/2005               I....S..............................N...NK............T..................................R...I.DP...
A/equine/Lincolnshire/1/2007              I....S..............................N...NK............T..................................R...I.DP...
A/equine/Beuvron-en-Auge/2/2009           I....S..............................N...NK............T..................................R...I.DP...
A/equine/Belfond/6-2/2009                 I....S..............................N...NK............T..................................R...I.DP...
A/equine/Florida/146609/2011              I....S..............................N...NK............T..................................R...INDP...
A/equine/Malaysia/1/2015                  I....S..............................N...NK............T..................................R...I.DP...
A/equine/Paris/1/2018                     I....S..............................N...NK............T..................................R...I.DP...
A/Equus  ferus caballus/USA/51053/2019    I....S..............................N...NK............T..................................R...I.DP...
A/donkey/Nigeria/VRD19/ZM20 T10 19RS459-5/2019 I....S..............................N...NK............T..................................R...I.DQ...
```

Supp Fig 4 KLEIJ et al., 2023

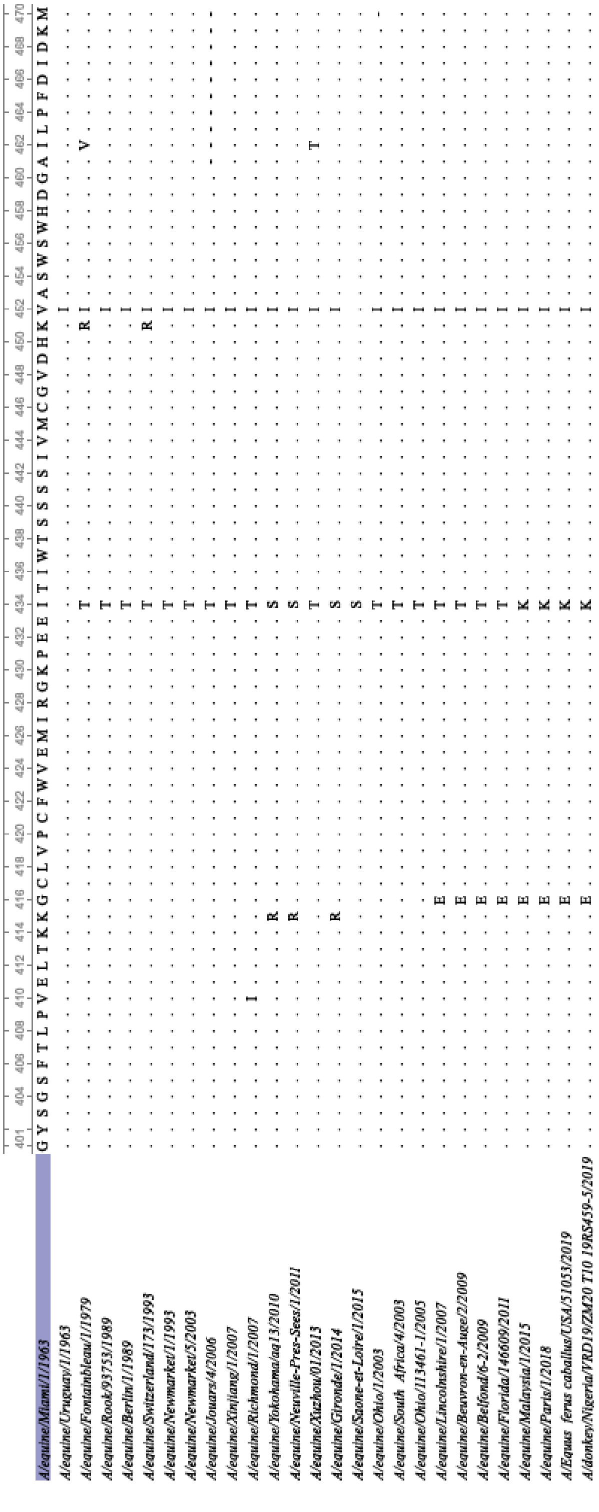


**Supp Fig 4 KLEIJ et al., 2023**

| strains | 1_PB2 | 2_PB1 | 3_PA | 4_HA | 5_NP | 6_NA | 7_MP | 8_NS |
|---|---|---|---|---|---|---|---|---|
| A/equine/Miami/1/1963 | CY028843 | CY028842 | CY028841 | CY028836 | CY028839 | L06580 | CY028837 | CY028840 |
| A/equine/Uruguay/1/1963 | CY032428 | CY032427 | CY032426 | CY032421 | CY032424 | CY032423 | CY032422 | CY032425 |
| A/equine/Fontainbleau/1/1979 | CY032412 | CY032411 | CY032410 | CY032405 | CY032408 | CY032407 | CY032406 | CY032409 |
| A/equine/Berlin/1/1989 | CY032420 | CY032419 | CY032418 | CY032413 | CY032416 | CY032415 | CY032414 | CY032417 |
| A/equine/Rook/93753/1989 | CY032332 | CY032331 | CY032330 | CY032325 | CY032328 | CY032327 | CY032326 | CY032329 |
| A/equine/Switzerland/173/1993 | CY032364 | CY032363 | CY032362 | CY032357 | CY032360 | CY032359 | CY032358 | CY032361 |
| A/equine/Newmarket/1/1993 | FJ375218 | FJ375230 | FJ375226 | FN399025 | FJ375214 | FJ375222 | FJ375234 | FJ375210 |
| A/equine/Ohio/1/2003 | DQ124177 | DQ124172 | DQ124187 | DQ124192 | DQ124184 | DQ124168 | DQ124188 | DQ124186 |
| A/equine/Newmarket/5/2003 | FJ375221 | FJ375233 | FJ375228 | FJ375213 | FJ375216 | FJ375224 | FJ375236 | FJ375209 |
| A/equine/South_Africa/4/2003 | ON797667 | ON797668 | ON797669 | ON797670 | ON797671 | ON797672 | ON797673 | ON797674 |
| A/equine/Ohio/113461-1/2005 | CY067315 | CY067316 | CY067317 | CY067318 | CY067319 | CY067320 | CY067321 | CY067322 |
| A/equine/Jouars/4/2006 | KY241366 | KY241372 | KY241378 | JX091752 | KY241321 | KY241348 | KY241354 | KY241360 |
| A/equine/Xinjiang/1/2007 | EU794540 | EU794541 | EU794542 | EU794543 | EU794544 | EU794545 | EU794546 | EU794547 |
| A/equine/Richmond/1/2007 | KF559332 | KF559333 | KF559334 | FJ195395 | KF559335 | KF559336 | KF559337 | FJ195429 |
| A/equine/Lincolnshire/1/2007 | KF559338 | KF559339 | KF559340 | FJ195398 | KF559341 | KF559342 | KF559343 | FJ195427 |
| A/equine/Belfond/6-2/2009 | KY241367 | KY241373 | KY241379 | KY241314 | KY272866 | KY241350 | KY241355 | KY241361 |
| A/equine/Yokohama/aq13/2010 | LC369077 | LC369078 | LC369079 | LC369080 | LC369081 | LC369082 | LC369083 | LC369084 |
| A/equine/Neuville-Pres-Sees/1/2011 | KY241368 | KY241374 | KY241380 | KY241316 | KY241322 | KY241347 | KY241359 | KY241362 |
| A/equine/Florida/146609/2011 | MF173363 | MF173219 | MF173307 | MF173407 | MF173304 | MF173237 | MF173292 | MF173196 |
| A/equine/Xuzhou/01/2013 | KF806992 | KF806991 | KF806990 | KF806985 | KF806988 | KF806987 | KF806986 | KF806989 |
| A/equine/Gironde/1/2014 | KY241370 | KY241376 | KY241382 | KY241319 | KY241324 | KY241352 | KY241357 | KY241364 |
| A/equine/Saone-et-Loire/1/2015 | KY241371 | KY241377 | KY241383 | KY241320 | KY241325 | KY241353 | KY241358 | KY241365 |
| A/equine/Malaysia/1/2015 | MH135220 | MH135221 | MH135222 | MH135223 | MH135224 | MH135225 | MH135226 | MH135227 |
| A/Equus_ferus_caballus/USA/51053/2019 | MT212122 | MT212123 | MT212124 | MT212125 | MT212126 | MT212127 | MT212128 | MT212129 |
| A/donkey/Nigeria/VRD19_ZM20_T10_19RS459-5/2019 | MT272441 | MT272440 | MT272439 | | MT272437 | MT272436 | MT272435 | MT272438 |
| A/donkey/Senegal/44/2019 | | | | MT272434 | | | | |

| | HA | | | | | | | | | | | | | | | | | | | | |
|---|---|---|---|---|---|---|---|---|---|---|---|---|---|---|---|---|---|---|---|---|---|
| Amino acid position | 6 | 7 | 47 | 62 | 63 | 78 | 104 | 138 | 159 | 163 | 188 | 223 | 272 | 291 | 372 | 379 | 450 | 454 | 474 | 495 | 527 |
| A/equine/Ohio/1/2003 | S | G | S | R | N | A | D | A | S | T | N | V | V | D | A | G | K | Q | D | A | I |
| A/equine/South Africa/4/2003 | - | - | - | - | - | - | - | - | - | - | - | - | - | - | - | - | - | - | - | V | - |
| A/equine/Ohio/113461-1/2005 | - | - | - | - | - | - | - | - | - | - | - | - | - | - | - | - | - | - | - | - | - |
| A/equine/Richmond/2007 | - | N | - | - | - | V | - | - | N | - | - | - | - | E | T | E | - | - | - | - | - |
| A/equine/Beuvron-en-Auge/2/2009 | - | D | - | K | - | - | N | S | - | - | - | I | A | - | - | - | - | - | - | - | V |
| A/equine/Paris/1/2018 | N | D | P | K | D | - | N | S | - | I | T | I | - | - | T | - | R | L | N | - | - |

| | NA | | | | | | | | | | | | | | | | | | | | | | | | | | |
|---|---|---|---|---|---|---|---|---|---|---|---|---|---|---|---|---|---|---|---|---|---|---|---|---|---|---|---|
| Amino acid position | 8 | 9 | 12 | 13 | 21 | 35 | 40 | 42 | 47 | 66 | 68 | 70 | 74 | 76 | 78 | 147 | 186 | 191 | 203 | 235 | 252 | 258 | 260 | 271 | 337 | 410 | 416 | 434 |
| A/equine/Ohio/1/2003 | I | A | F | T | S | A | R | D | E | Y | I | T | M | K | S | I | K | V | V | D | K | N | K | G | N | V | E | K |
| A/equine/South Africa/4/2003 | - | T | S | A | N | V | E | G | G | H | T | A | I | R | P | V | - | I | - | N | R | D | R | - | S | I | G | T |
| A/equine/Ohio/113461-1/2005 | M | - | - | A | N | - | - | - | G | - | T | - | I | R | - | V | R | - | - | - | R | D | R | | - | - | G | T |
| A/equine/Richmond/2007 | - | - | - | A | N | V | - | - | G | - | T | - | I | R | - | V | - | - | - | - | R | D | R | E | S | - | G | T |
| A/equine/Beuvron-en-Auge/2/2009 | - | - | - | A | N | V | - | - | G | - | T | - | I | R | - | V | - | - | - | - | R | D | R | - | S | - | G | T |
| A/equine/Paris/1/2018 | M | - | - | A | N | - | - | - | G | - | T | - | I | R | - | V | - | - | I | - | R | D | - | - | - | - | - | T |

**Supp Table 2 KLEIJ et al., 202**

| Genomic segment | Primer Name | Primer sequence (5’-> 3’) | Amplicon size (bp) |
|---|---|---|---|
| segment 1 | sgt1-PCR-U12 | AGCGAAAGCAGGTCAAATATATTC | 2341 |
| | sgt1-PCR-U13 | AGTAGAAACAAGGTCGTTTTTAAAC | |
| segment 2 | sgt2-PCR-U12 | AGCGAAAGCAGGCAAACCATTTG | 2341 |
| | sgt2-PCR-U13 | AGTAGAAACAAGGCATTTTTTCATG | |
| segment 3 | sgt3-PCR-U12 | AGCGAAAGCAGGTACTGATCCA | 2233 |
| | sgt3-PCR-U13 | AGTAGAAACAAGGTACTTTTTTGGA | |
| segment 4 | sgt4-PCR-U12 | AGCAAAAGCAGGGGATATTTCTGT | 1768 |
| | sgt4-PCR-U13 | AGTAGAAACAAGGGTGTTTTTAACT | |
| segment 5 | sgt5-PCR-U12 | AGCAAAAGCAGGGTAGATAATCA | 1565 |
| | sgt5-PCR-U13 | AGTAGAAACAAGGGTATTTTTCTTT | |
| segment 6 | sgt6-PCR-U12 | AGCAAAAGCAGGAGTTTAAAATGAA | 1460 |
| | sgt6-PCR-U13 | AGTAGAAACAAGGAGTTTTTTCGT | |
| segment 7 | sgt7-PCR-U12 | AGCAAAAGCAGGTAGATATTTAAAG | 1027 |
| | sgt7-PCR-U13 | AGTAGAAACAAGGTAGTTTTTTAC | |
| segment 8 | sgt8-PCR-U12 | AGCAAAAGCAGGGTGACAAAAAC | 890 |
| | sgt8-PCR-U13 | AGTAGAAACAAGGGTGTTTTTTATC | |

**Supp Table 3 KLEIJ et al., 202[illegible]**
